# Gait Adaptations in Step Length and Push-off Force during Walking with Functional Asymmetry

**DOI:** 10.1101/2025.01.05.631404

**Authors:** Tomislav Baček, Yufan Xu, Liuhua Peng, Denny Oetomo, Ying Tan

## Abstract

Human walking is remarkably adaptable, allowing individuals to maintain efficiency and stability across diverse conditions. This study investigates how healthy young adults adapt step length and push-off force to varying speeds and cadences under functional asymmetry induced by a unilateral knee constraint, simulating hemiparetic gait. A dataset of 19 participants walking across 30 conditions was used to examine these adaptations in both absolute terms and symmetry metrics. Results reveal that functional asymmetry disproportionately impacts propulsion, with constrained-leg force decreasing significantly at higher speeds. Step length symmetry remains stable, suggesting a prioritisation of spatial over kinetic symmetry, likely to optimise walking energetics and maintain anterior-posterior balance. Statistical models demonstrated good within-dataset performance but limited generalisability in cross-dataset predictions, where differences in population cohorts and experimental designs reduced accuracy. These findings provide insights into the mechanisms driving gait adaptations and highlight critical limitations in applying statistical models to patient populations, underscoring the importance of representative datasets and caution when extrapolating findings across populations.

## 1 Introduction

Human walking is remarkably versatile, driven by complex adaptation mechanisms that allow individuals to maintain efficiency, stability, and control in a variety of conditions. Healthy individuals generally prefer to walk at a speed that minimises energy expenditure per unit of distance traveled [1, 2]. To achieve this, they actively adjust gait variables, including step width [3], step length [4], step time [5], and push-off force [6], while avoiding penalties associated with asymmetries [5]. These adjustments vary with walking speed: changes in step width support lateral balance [7, 8], while changes in step length accommodate the increased demands of push-off [9, 10], all while maintaining a sufficient margin of stability [11]. This adaptability allows people to seamlessly walk on slopes [12, 13], in the presence of lateral perturbations [8, 14], or with physical changes of aging [15, 16].

In contrast, hemiparetic individuals walk with slower speeds [17, 18] and higher energetic cost [19], exhibiting spatio-temporal asymmetries, such as the extended stance phase on the non-paretic side, prolonged swing phase on the paretic side [20], and reduced paretic propulsion [21]. These asymmetries manifest in both spatio-temporal asymmetry [22–24] and asymmetries in push-off force and impulse [25, 26], with some patients taking longer paretic steps while others take shorter ones [20]. However, despite these differences, hemiparetic individuals can still independently adjust gait parameters, relying on strategies that include increased non-paretic propulsion or wider steps [27, 28] to maintain stability and forward progression.

While much is known about gait adaptations in both healthy and patients, significant gaps remain. Many studies rely on speed-matching or age-matching between healthy and patient groups, which fail to account for intrinsic biomechanical differences [29, 30]. Such approaches can mask critical insights by overlooking how baseline differences – such as muscle weakness, pain, or comorbidities – alter gait dynamics independently of age or walking speed. Moreover, walking speed alone is a limited indicator of gait function and recovery, providing an incomplete picture of metrics like step length and forward propulsion [31]. These limitations challenge the validity of using matched-group analyses or statistical models trained on healthy populations to explain or predict patient gait, as such models tend to overfit dataset-specific dynamics.

To address these gaps, this study employs a unique dataset of healthy individuals walking with a sagittal-plane functional asymmetry induced using unilateral knee constraint [32]. This within-subject design, where each participant serves as their own control, isolates the effects of asymmetry on step length and push-off force (i.e., propulsion) mechanisms, allowing for comparisons unaffected by the confounding factors common in patient populations. By systematically varying speed and cadence, the study explores the interplay between these factors and symmetry metrics under both symmetrical (*free*) and asymmetrical (*constrained*) conditions.

Step length and push-off force are key to understanding anterior-posterior balance [8] and the metabolic and mechanical costs of walking [9]. These variables are increasingly used as biofeedback targets in hemiparetic rehabilitation [10], making it crucial to understand their adaptability across conditions. This study leverages statistical models not only to analyse gait adaptations but also to critically evaluate the limitations of such models when applied across datasets. In doing so, the study aims to:

- investigate the mechanisms underlying step length and forward propulsion adaptations to speed and cadence in the presence of functional asymmetry;
- assess the utility and generalisability of statistical models, demonstrating how dataset-specific dynamics limit their applicability to clinical populations.

## 2 Materials and methods

### 2.1 Walking dataset

We used publicly available dataset from [32], hereafter referred to as B24, to investigate the effects of functional gait asymmetry on step length (SL) and peak push-off force (PO). In this context, functional asymmetry refers to a constrained condition in which a knee brace was used to unilaterally restrict knee joint movement. The dataset comprises data from 21 neurotypical young adults (age 30*±*8 years, body mass 72.7*±*12.3 kg, height 1.72*±*0.09 m), all of whom were free of any lower-extremity injury and wore low-profile shoes during the trials. Two participants (Subjects 7 and 12 in the original dataset) only completed the first data collection session and were excluded from this analysis. No data were missing for the remaining 19 participants analysed in this paper. The anthropometric characteristics of participants in the B24 dataset are given in the Appendix (Table A1). The ethics committee of The University of Melbourne approved the study in [32] (reference number: 2021-20623-13486-3).

The trials were conducted on a dual-belt instrumented treadmill, with each lasting 5 minutes. Participants completed a total of 30 trials: 15 without any constraints (hereafter referred to as *Free*) and 15 with their left knee joint fully extended via a passive knee brace (hereafter referred to as *Constrained*). In both conditions, participants walked at three speeds (0.4, 0.8, and 1.1 m/s) and five step frequencies peer speed, guided by a metronome (90%, 95%, 100%, 110%, and 120% of the preferred cadence, presented in random order). The 30 trials took place across two sessions on separate days, with 15 trials per day. The five cadences at each speed were organised into a continuous 25-minute walking bout with no breaks between cadences. Bouts were arranged such that no two consecutive bouts involved the same walking conditions (either *Free* or *Constrained*) within a session, and all three speeds were included in each session. Marker data were collected at 100 Hz and ground reaction force (GRF) data at 1000 Hz. A detailed description of the study is provided in [32].

For this analysis, we separated the B24 dataset into four groups, corresponding to the left and right legs in each of the two conditions (*Free* and *Constrained*). We refer to the data from the left leg during unconstrained walking as *Free Left* and the data from the right leg as *Free Right*. Similarly, *Constrained Left* represents data from the left leg during constrained walking and *Constrained Right* refers to the data from the right leg during constrained walking. Note that it is always the left leg that is constrained during all *Constrained* walking in the experiment. Hence, *Constrained Right* is the data of the unconstrained right leg in the *Constrained* walking condition. Given the strong correlation of gait parameters between *Free Left* and *Free Right*, we use *Free Left* as the baseline for the *Free* condition.

### 2.2 Data processing and analysis

All data processing – including filtering, segmenting, and grouping – was performed using custom-written scripts in Matlab 2024a. Raw GRF data were filtered using a low-pass Butterworth filter with a 6 Hz cut-off frequency [33]. The vertical component of the GRF signal was used to segment data into gait cycles, with a threshold set at 5% of the peak amplitude (e.g., for a 75 kg person, the threshold would be 75*→*9.81*→*0.05 = 37 N). In our analysis, the same number of cycles was used for both legs, and all gait metrics represent an average over the last 60 seconds of each 5-minute test.

Statistical modelling was done in Python 3.10.10 using *statsmodels*, *scipy*, and *numpy* packages, with a Linear Mixed Models (LMM) approach [34]. Each participant’s data was treated as a distinct group to account for constant anthropometric variables within each group while allowing variations in their gait metrics. Data from the left and right legs were modelled separately, treating peak push-off force (PO) and step length (SL) as response variables (model outputs), and walking speed, its square, cadence (for PO) or trailing limb angle (TA, for SL), and anthropometric data (sex, age, weight, leg length) as explanatory variables (model inputs). Both PO and SL models include a fixed effect for speed given the known relationship between the speed and the two response variables [35]. Models also include quadratic relationship with speed, similar to the models in [10], and a fixed intercept. In addition to these fixed effects, both models include a random slope for speed to account for the repeated measures structure of the B24 dataset and variability across participants. We report the best performing PO and SL models with speed variance, reflecting the average individual effect of speed.

Model estimation quality across combinations of independent variables was assessed using the Akaike Information Criterion (AIC), and we report AIC values for all models, along with the parameters for the best model for each model output. We evaluate the predictive model performance using several complementary metrics, including R-Squared (R^2^), Mean Absolute Error (MAE), and Mean Absolute Percentage Error (MAPE)^1^. The accuracy of the predictive model was evaluated on the B24 dataset and further tested on an independent, publicly available dataset by [36] (details in Appendix).

Statistical data analyses were conducted in Python 3.10.10 using *scipy* package. The effects of speed, cadence, and condition (free vs. constrained) on gait variables were assessed using two-way repeated measures ANOVA (RMANOVA) at a significance level of *p*=0.05. Where statistically significant differences were found, pairwise post-hoc analyses were carried out using Tukey’s honestly significant difference (HSD) test, with Bonferroni corrections applied. We categorise statistical significance as follows: weak significance (0.01*<p↑*0.05), moderate significance (0.001*<p↑*0.01), and strong significance (*p↑*0.001).

### 2.3 Gait metrics

A gait cycle is defined as the time between two successive heel strikes of the same leg. Peak push-off force (PO) represents the maximum amplitude of the fore-aft component of the GRF during the stance phase, which spans from heel strike to toe-off of the same leg. Step length (SL) is the fore-aft distance between the two calcaneous (heel) markers at the time of the leading leg’s heel strike. Symmetry is defined as the percentage ratio between the two legs: left vs. right for PO (100% indicating symmetry) and left vs. left plus right for SL (50% indicating symmetry).

Trailing limb angle (TA) is defined as the maximum hip extension angle. The hip joint angle trajectory was calculated following the methods outlined in Research Methods in Biomechanics [37] and according to International Society of Biomechanics (ISB) guidelines in [38]. Leg length was measured as the distance from the anterior superior iliac spine (ASIS) to the ipsilateral medial malleolus during static calibration in a standing position.

## 3 Results

### 3.1 Statistical modelling (absolute gait metrics)

#### 3.1.1 Model estimation quality

We trained models for peak push-off force (PO) and step length (SL) using combinations of walking speed and its square (*v* and *v*^2^) along with anthropometric variables (Age, Sex, Weight, and Leg Length). Additionally, we included step frequency (*f*) for the PO model and trailing limb angle (TA) for the SL model (using speed and cadence as SL model inputs would not be an estimate but a closed form solution due to *SL* = *v/f*). We define three model types: *Model1*, trained with *v* and *v*^2^ only; *Model2*, trained with *v*, *v*^2^, and either *f* or TA; and *Model3*, trained with *v*, *v*^2^, either *f* or TA, and all four anthropometric variables. Models incorporating individual anthropometric variable are not presented here, as analysis indicated that no single variable consistently achieved significance across multiple models.

Table 1 provides Aikake Information Criterion (AIC) values for both model outputs (PO, SL), including all three model levels and four data groups. As the table shows, walking speed and either cadence (for PO model) or TA (for SL model) were the only model parameters that significantly improved model estimation quality, while adding anthropometric variables as model inputs had minimal to no impact. For this reason, all further modelling of PO and SL is performed using *Model2*. Table 2 presents the parameters for the best overall model (*Model3*), specifically for the *Free Right* condition in the case of PO and the *Constrained Left* condition for SL. A more detailed analysis of model estimation quality can be found in the Appendix.

**Table 1:** AIC values for PO and SL model estimation.

|  | Push-off force (PO) |  |  | Step length (SL) |  |  |
| --- | --- | --- | --- | --- | --- | --- |
| Data group | Model1 | Model2 | Model3 | Model1 | Model2 | Model3 |
| Free Left | -162.18 | -260.44 | -270.64 | -782.61 | -1008.58 | -1036.79 |
| Free Right | -192.03 | -270.77 | -283.26 | -752.03 | -958.57 | -985.08 |
| Constrained Left | -13.265 | -94.35 | -94.11 | -709.44 | -1036.96 | -1055.63 |
| Constrained Right | -93.86 | -175.76 | -182.54 | -746.34 | -853.99 | -870.60 |

**Table 2:** Parameters for the model that best fit PO and SL overall (*Model3*).

|  | intercept | v | v<sup>2</sup> | cadence / TLA | leg length | sex | age | weight | v variance |
| --- | --- | --- | --- | --- | --- | --- | --- | --- | --- |
| PO | 0.84 | 1.668 | 0.338 | -0.008 | -0.417 | 0.036 | 0.006 | -0.002 | 0.026 |
| SL | -0.197 | 0.74 | -0.332 | -0.012 | 0.201 | 0.03 | 0.003 | -0.001 | 0.004 |

Cadence only used in PO model, and TLA only in SL model.

#### 3.1.2 Model prediction quality

Fig. 1 illustrates the prediction of PO using a model trained on the *Free Left* data (left leg during free walking). The model accurately predicts PO of the contralateral (right) leg in free walking across all three speeds (*Free Right*; Fig. 1, left), achieving a Mean Absolute Error (MAE) of 0.16 N/kg and a Mean Absolute Percentage Error (MAPE) of 14.3%. However, prediction accuracy decreases when applied to constrained walking (*Constrained Right*; Fig. 1, right), particularly at higher walking speeds, where the model sometimes underestimates the measured force. This is reflected in lower R^2^ and higher errors (MAE = 0.21 N/kg and MAPE = 15.3%) compared to the *Free* condition. Notably, the right leg remains unconstrained in both cases, showing minimal change in PO on the right leg even when the left leg is constrained. In all models below, each datapoint represents one test per participant.

**Figure 1:**
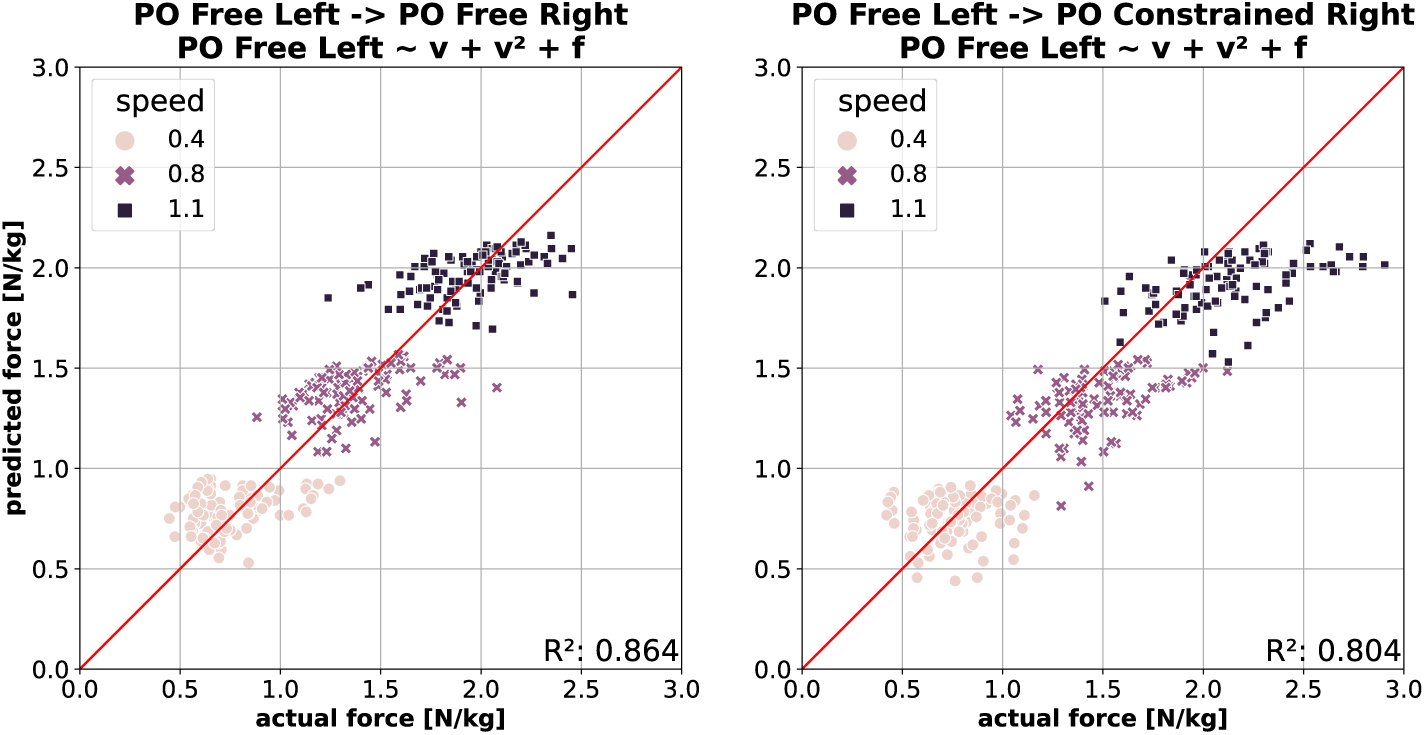
PO prediction using model trained on the free walking data. *(**Left**)* Predicting contralateral leg in free walking. *(**Right**)* Predicting contralateral leg in constrained walking.

Fig. 2 shows the PO prediction across the two conditions (from free to constrained and vice versa). A model trained on the free walking data (*Free Left*) performs well in predicting PO in constrained walking (*Constrained Left*; Fig. 2, left) at slow speed, though its accuracy decreases as speed increases. At higher speeds, the model tends to overestimate measured force, which is reflected in lower R^2^ and higher error (MAE = 0.24 N/kg and MAPE = 23.5%). A similar, albeit less pronounced, trend appears when predicting free walking (*Free Left*) using a model trained on constrained walking data (*Constrained Left*; Fig. 2, right). Here, higher walking speeds lead underestimation of the measured push-off force, resulting in denser data distribution. This model performs better overall, as indicated by a higher R^2^ and lower errors (MAE = 0.22 N/kg and MAPE = 17.0%.).

**Figure 2:**
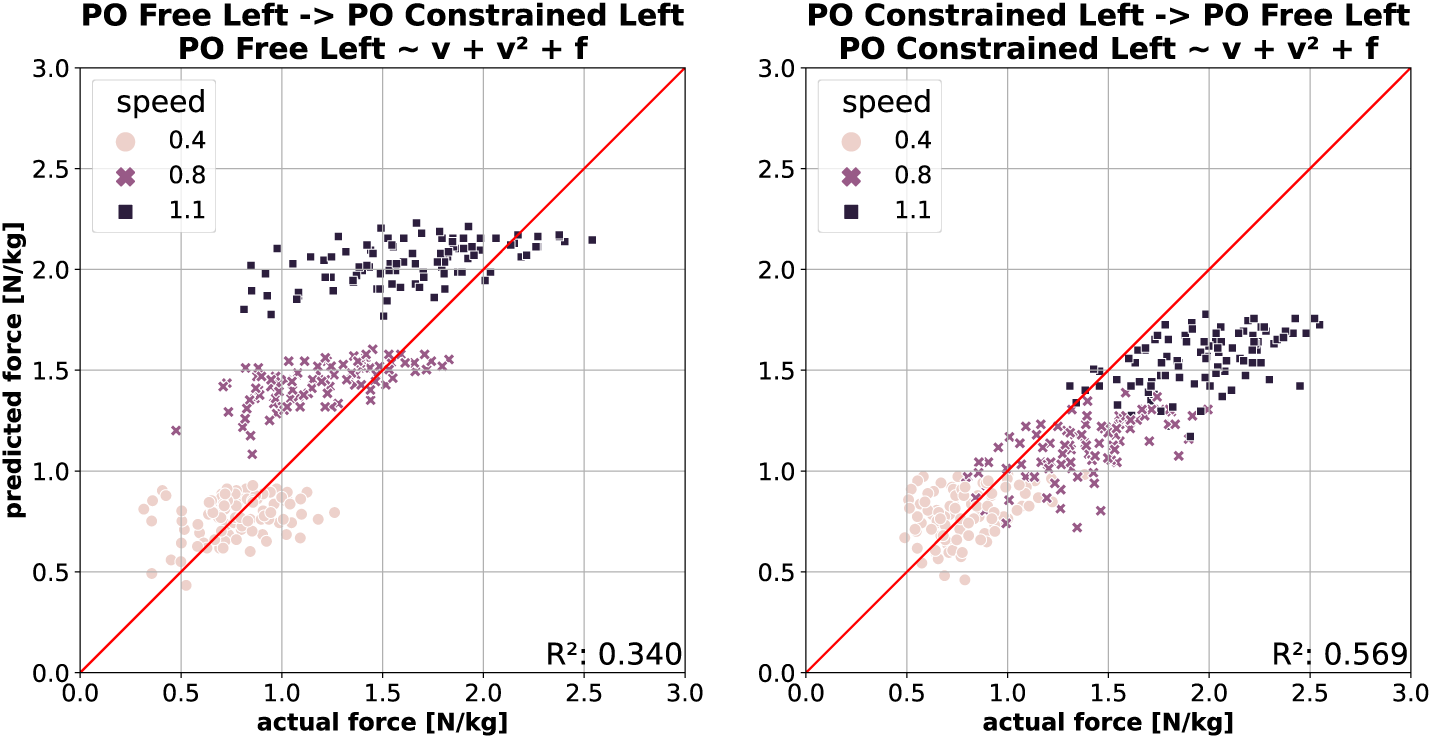
PO prediction across conditions. *(**Left**)* Predicting constrained left leg using model trained on the free left leg data. *(**Right**)* Predicting free left leg using model trained on the constrained left leg data.

Prediction of SL on the contralateral leg using a model trained on the free walking data (*Free Left*) is shown in Fig. 3. The model predicts SL with comparable accuracy in both free walking (*Free Right*; Fig. 3, left) and constrained walking (*Constrained Right*; Fig. 3, right), though with slightly reduced accuracy in the latter. The difference is reflected by lower R^2^ and higher errors in constrained walking (MAE = 0.047 N/kg and MAPE = 10.4% vs. MAE = 0.058 N/kg and MAPE = 13.2%). Unlike the PO prediction, walking speed does not appear to influence SL prediction accuracy. Note that the right leg remains unconstrained in both cases.

**Figure 3:**
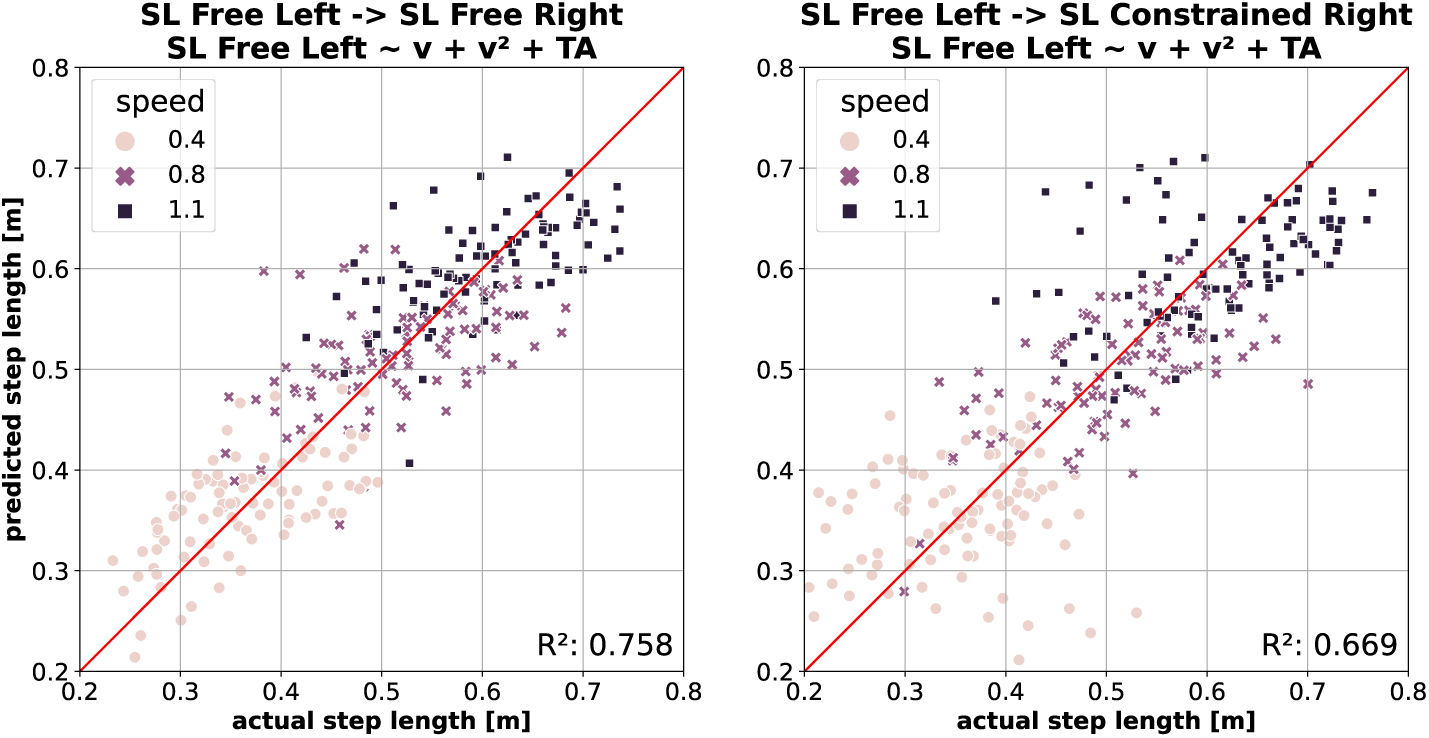
SL prediction using model trained on the free walking data. *(**Left**)* Predicting contralateral leg in free walking. *(**Right**)* Predicting contralateral leg in constrained walking.

Fig. 4 illustrates SL prediction across conditions. Predicting SL for the constrained leg (*Constrained Left*) using a model trained on free walking data (*Free Left*; Fig. 4, left) yields the best SL prediction performance, as reflected by the highest R^2^ and lowest errors (MAE = 0.044 and MAPE = 9.6%). Predicting SL in the opposite direction – using a model trained on constrained walking data (*Constrained Left*) to predict free walking (*Free Left*; Fig. 4, right) – is only slightly less accurate, with high R^2^ and low errors (MAE = 0.045 and MAPE = 9.7%). Similar to SL predictions within the free walking condition, walking speed does not significantly affect prediction quality in this context.

**Figure 4:**
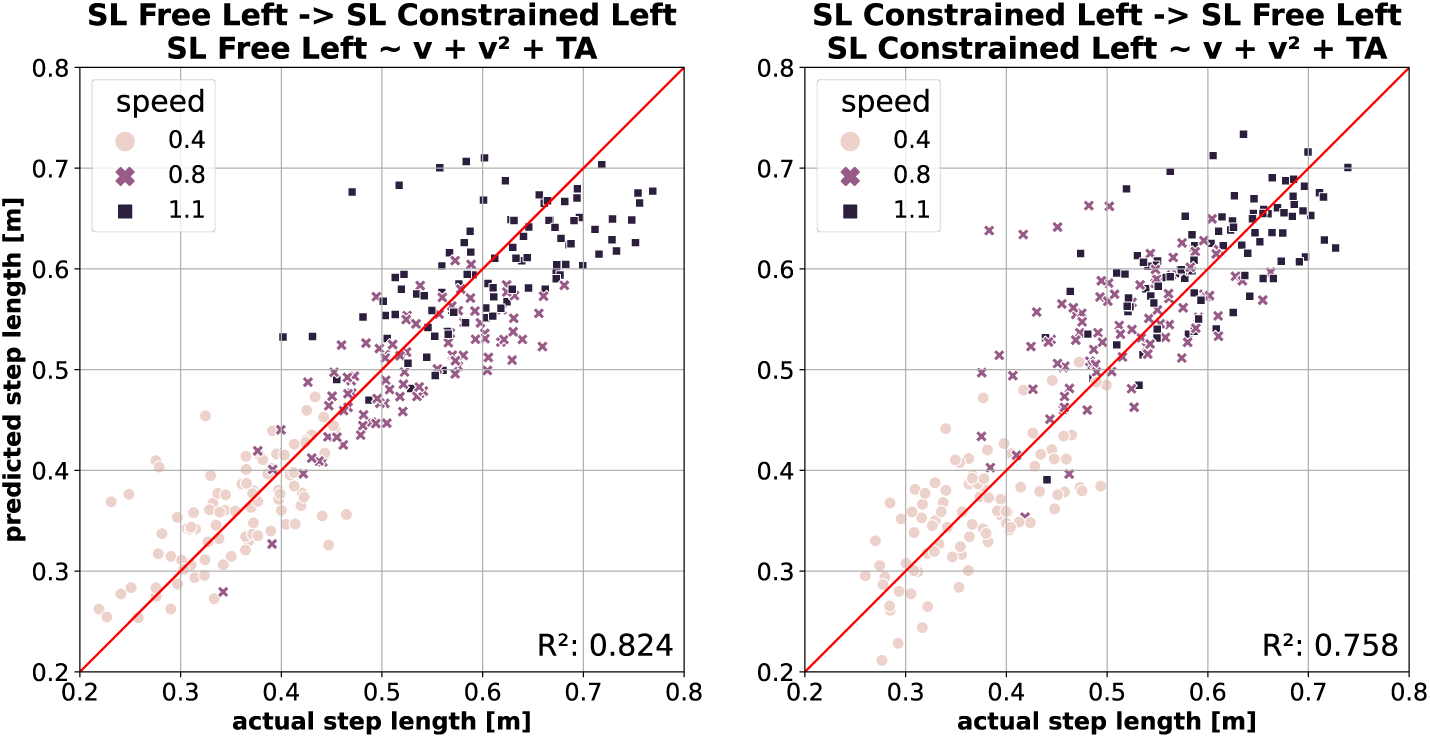
SL prediction across conditions. *(**Left**)* Predicting constrained left leg using model trained on the free left leg data. *(**Right**)* Predicting free left leg using model trained on the constrained left leg data.

### 3.2 Spatial and kinetic asymmetries (relative gait metrics)

#### 3.2.1 Peak push-off force (PO) symmetry

As shown in Fig. 1, the right leg generates peak push-off force (PO) consistent with predictions from a model trained on free walking, even when participants walked with constraints on their contralateral leg. Conversely, Fig. 2 shows that this symmetry does not hold for the left leg during constrained walking, where the peak push-off force (PO) is lower than predicted by a model trained on free walking. This discrepancy suggests a change in PO symmetry, particularly at higher walking speeds.

Fig. 5 (*top*) illustrates PO symmetry across conditions, supporting the observations discussed above. During free walking (blue bars), participants on average displayed strong symmetry in their propulsive forces at 0.8 m/s (101.2*±*11.7% across cadences; mean*±*standard deviation) and 1.1 m/s (100.9*±*7.5% across cadences), with slightly higher PO in their left leg at 0.4 m/s (109.4*±*14.9% across cadences). Cadence had no statistically significant effect on PO symmetry at any speed; however, there were weak statistically significant differences in PO symmetry across the speeds when averaging across cadences: *p*=0.03 between 1.1 and 0.4 m/s and *p*=0.04 between 0.8 and 0.4 m/s.

**Figure 5:**
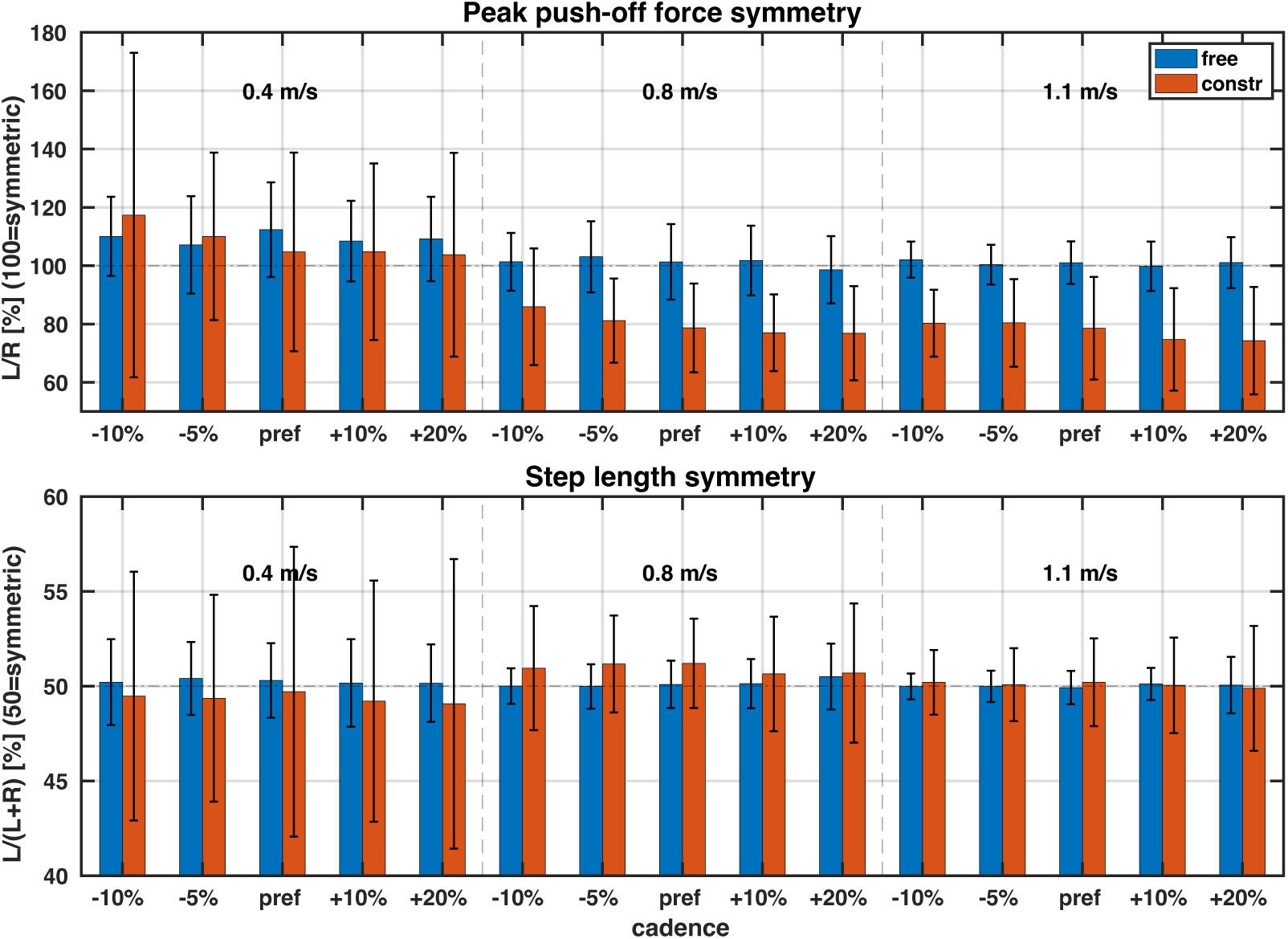
PO and SL symmetry. Data are average across all participants. Three walking speeds are separated by vertical dashed lines, and two conditions (free, constrained) are colour-coded. Horizontal dashed lines indicate perfect symmetry. Note different definitions of the symmetry between PO and SL. *(**Top**)* Peak push-off force symmetry (values below 100 indicate more force by the right leg). *(**Bottom**)* Step length symmetry (values below 50 indicate shorter left step).

Applying a knee brace on the left leg affected PO symmetry, particularly at 0.8 and 1.1 m/s. At the slowest speed (0.4 m/s), the constraint increased PO symmetry variability, although average PO symmetry remained similar to that during free walking (blue vs. red bars; *p>*0.99), with an average PO symmetry of 108.2*±*36.7% across cadences. At 0.8 and 1.1 m/s, PO symmetry on average decreased by approximately 20 percentage points compared to free walking, with the left leg producing less peak propulsive force; across cadences, PO symmetry averaged 79.9*±*15.8% at 0.8 m/s and 77.7*±*16.1% at 1.1 m/s. Similar to free walking, cadence had no statistically significant effect on PO symmetry at either of the two speeds. Furthermore, there were no statistically significant differences in PO symmetry between 0.8 and 1.1 m/s during constrained walking, although both speeds were statistically significantly different from 0.4 m/s (strong significance, *p<*0.001).

#### 3.2.2 Step length (SL) symmetry

The modelling of SL, depicted in Fig. 3 and 4, indicated there should be no major asymmetries in SL across speeds in either free or constrained walking. The SL symmetry visualisation in Fig. 5 (*bottom*) confirms this observation. During free walking (blue bars), participants on average walked with strong SL symmetry across all three speeds (50.3*±*2.1% at 0.4 m/s, 50.1*±*1.3% at 0.8 m/s, and 50.01*±*0.9% at 1.1 m/s, averaged across cadences), with cadence having no statistically significant effect on SL symmetry at any speed. Unlike PO symmetry, SL symmetry was unaffected by the knee constraint (*p>*0.59 across all comparisons; notably, the only statistically significant difference, albeit a weak one, was between 0.4 and 0.8 m/s in constrained walking: *p*=0.04). Although SL symmetry variability increased across all three speeds, particularly at 0.4 m/s, the overall symmetry remained stable. On average across participants and cadences, SL symmetry during constrained walking remained close to strong symmetry: 49.4*±*6.7% at 0.4 m/s, 50.2*±*2.9% at 0.8 m/s, and 50.1*±*2.4% at 1.1 m/s.

## 4 Discussion

In this study, we investigated how healthy young adults adapt their step length and propulsion in response to varying speeds and cadences during free and emulated hemiparetic walking. The latter was achieved through a passive knee brace worn by the study participants on their left leg. Using statistical models, we quantified these adaptations in absolute terms and assessed how well these models predict data from an independent, publicly available dataset. Additionally, we examined changes in step length and propulsion symmetries. Our findings provide a nuanced understanding of the mechanisms driving gait adaptations under functional asymmetry and establish a framework for comparing these adaptations across datasets.

### 4.1 Walking dataset

Human gait is often studied by comparing adaptations across different groups, including speed- or age-matching experimental designs [14, 16]. A major limitation of these approaches is their inability to account for biomechanical variations unrelated to walking speed or age, a challenge that is particularly true in patient populations [39]. The dataset used in this study – from [32] – is unique in its within-subject experimental design, which isolates gait adaptations caused specifically by functional asymmetry from external, confounding factors (e.g., muscle weakness).

To achieve this, participants walked with a unilateral passive knee brace that restricted left knee flexion, consequently limiting their ankle plantarflexion angle during push-off and prompting compensatory gait adjustments similar to those seen in hemiparetic patients – see [40]. This experimental design anticipated that increasing walking speed would present as a more demanding task due to naturally higher joint mobility demands at faster walking speeds [41], an expectation validated by the results here. Although a metronome guided their step frequency, participants were allowed to make small adjustments to step length (SL), step time (ST), and their respective symmetries, provided that their overall gait speed, determined by *stride* length and *stride* time, matched the treadmill’s imposed speed.

### 4.2 Statistical modelling of step length (SL) and peak push-off force (PO)

We started by developing models to predict the amplitudes (i.e., absolute values) of peak push-off force (PO; a component of the GRF in the anterior-posterior direction) and step length (SL) using gait speed and anthropometric variables as model inputs. Additionally, we included cadence as the PO model input and trailing limb angle (TA) as the SL model input. Previous studies have built similar models for SL [42], SL and PO [10], or alternatively, for lower limb joint angles and moments [43–45]. However, these studies have primarily focused on free walking, aiming to predict specific gait variables or develop biofeedback targets for clinical applications, often as alternatives to age- and sex-based normative values [46, 47]. In contrast, our study explored the effects of functional asymmetry on SL and PO and investigated whether these two variables adapt differently across a range of walking speeds.

During free walking, the ability of the models to estimate gait data across varying walking speeds was influenced only by speed and speed square, with anthropometric variables contributing minimally to this (see Table 1). This finding was consistent for both SL and PO and aligns with results reported by [10] – a comparison to [42] was not possible as their modelling was limited to a fixed speed. Due to the unique design of the dataset used, which includes five cadences per speed, our models also benefited from additional inputs – cadence for PO and trailing limb angle for SL – which allowed models to account for the enforced variations in SL (and consequently, PO) at the same walking speeds.

The models trained on free walking (*Free Left*) demonstrated high accuracy in predicting the contralateral (*Free Right*) leg’s PO (Fig. 1, left) and SL (Fig. 3, left), suggesting that both legs exhibit similar behaviours at a given speed and cadence. Notably, the same models, trained on free walking data (*Free Left*), were also able to predict contralateral leg PO (Fig. 1, right) and SL (Fig. 3, right) during constrained walking (i.e., *Constrained Right*) with only a slight reduction in accuracy, as measured by *R*^2^, MAE, and MAPE.

This is interesting since the predicted (right) leg remains unconstrained during walking with a knee brace (*Constrained Right*), suggesting that the right leg is only minimally influenced by the constraint applied to the left leg. The reduction in prediction accuracy is most pronounced for PO at higher speeds, where the model underestimates the actual PO. This indicates that increasing walking speed presents greater challenges for participants, prompting them to rely more on their unconstrained leg. A similar shift toward favouring the *unaffected* leg is also observed in hemiparetic patients, although it occurs independently of walking speed [31]. In contrast, the decrease in SL prediction accuracy does not appear to be speed-dependent. Instead, increased data dispersion across all three speeds contributes to the overall reduction in prediction quality. Interestingly, our findings reveal lower prediction accuracy for SL compared to PO, consistent with the results reported by [10], despite their model evaluations being conducted on an independent walking dataset.

We also evaluated model prediction accuracy for PO and SL of the left leg between conditions – from free to constrained walking, and vice versa. As shown in Fig. 2 (left), the model trained on free walking (*Free Left*) accurately predicts PO for the constrained leg (*Constrained Left*) at slow speeds. However, as walking speed increases, the model progressively overestimates the propulsive force of the constrained (left) leg. This indicates that participants generate less PO at higher speeds than what the free walking model predicts. Given that the data of the same participants were used to build the free walking model, it is the mechanics of the constraint – (close to fully) extended knee joint – and not musculoskeletal condition, such as muscle weakness or lack of muscle control, that cause a decrease in peak PO production.

This adaptation resembles, though is not necessarily identical to, the concept of a propulsive reserve observed in both elderly individuals [12] and hemiparetic patients [25] at their preferred walking speeds. It is plausible that all three groups – hemiparetic patients, elderly individuals, and healthy young adults walking with functional asymmetry (as in this study) – adopt a strategy of walking with lower-than-available propulsive forces, which may be a preferred compensatory mechanism in response to impairment, aging, or imposed constraints.

Unlike PO, SL prediction remains unaffected by the addition of the knee constraint. As Fig. 4 shows, using a model trained on free walking to predict constrained walking (left subfigure) and vice versa (right subfigure) yields similarly high accuracy. Similar to predictions on the unconstrained leg, walking speed has no discernible impact on SL prediction accuracy as the left leg in both the free (*Free Left*) and constrained walking (*Constrained Left*) exhibits similar SL. This is unexpected given the critical role that SL modulation plays in maintaining gait stability in the sagittal plane [8, 11]; as we’ve shown in [48], adaptations in step width are not the source of this.

The fact that the knee constraint does not influence the absolute value of SL on the constrained leg compared to the free leg suggests that participants did not rely on SL modulation to adjust their stability in the plane of walking. This behaviour might reflect a prioritisation of walking energetics over stability. As walking speed increases, humans typically increase their step length and step frequency concurrently [49] to avoid the metabolic penalties associated with disproportionately increasing one over the other [9]. Any potential reduction in sagittal-plane stability that might have happened as a consequence could indicate that participants either adopted alternative strategies for modulating fore-aft stability [6] or that the decrease in stability remained within tolerable limits, which aligns with the notion that humans prioritise sufficient rather than maximal gait stability [11].

### 4.3 Spatial (SL) and kinetic (PO) gait symmetries

Healthy young adults typically exhibit kinetic symmetry at walking speeds below 1.5 m/s. This includes symmetry in vertical, braking, and propulsive GRF components [50], as well as in propulsive and vertical impulses [51]. Our findings align with this: during free walking, participants on average displayed strong symmetry in peak push-off force at 0.8 and 1.1 m/s, with a slightly higher force on the left leg at 0.4 m/s (Fig. 5). Notably, this symmetry persisted across varying step frequencies, even during non-preferred stride lengths and times (i.e., 90, 95, 110, and 120% of preferred cadence). Similarly, our participants exhibited symmetry in forward (propulsive) and vertical (weight support) impulses, calculated as the ratio of left to right impulses (100% = symmetric). During free walking, weight bearing impulse symmetry demonstrated a slight tendency toward higher right-leg impulses, with an average of 97.65*±*2.4% across all speeds (see Fig. A1 in Appendix). Conversely, propulsive impulse symmetry showed a consistent tendency toward slightly higher left-leg impulses, with an average of 102.57*±*14.85% across all speeds (see Fig. A2 in Appendix). Despite the slight asymmetries, these results reinforce the broader notion that healthy adults generally maintain strong kinetic symmetry across a range of walking speeds.

Young healthy adults also generally exhibit strong symmetry in spatial and temporal step parameters. As walking speed increases, step length (SL) and step time (ST) increase concurrently, likely to mitigate the metabolic costs associated with disproportionately increasing only one [9]. In the absence of musculoskeletal injuries or impairments, humans tend to select an optimal combination of *stride* time and *stride* length at any given speed to minimise metabolic cost [1]. This is achieved by maintaining equal SLs and STs across both legs, thus avoiding metabolic penalties associated with step asymmetry [5, 52]. Consistent with findings on kinetic symmetries, our results also align with the literature on spatial symmetries: during free walking, participants exhibited strong SL symmetry across all speeds and cadences (Fig. 5, bottom). Following the symmetry criteria outlined in [53], where SL is considered symmetric if it falls within the range of 46.5% to 53.5% using the same symmetry definition as in this study, all participants but one (at 0.4 m/s) demonstrated symmetric SL (see Fig. A3 (left) in Appendix). Similarly, participants also exhibited strong ST symmetry across speeds and conditions^2^, which suggests that during free walking, participants maintained reduction of metabolic cost of walking as one of the priorities of gait.

During constrained walking, significant changes in peak push-off (PO) symmetry were observed. At 0.4 m/s, PO symmetry exhibited greater inter-person variability across all cadences. Notably, three participants changed from longer left step (PO symmetry *>*100%) to shorter left step (PO symmetry *<*100%), while two showed the opposite shift. At 0.8 and 1.1 m/s, PO symmetry significantly decreased, with nearly all participants showing reduced left-leg propulsion. For instance, at 0.8 m/s, of the nine participants with higher left peak push-off force during free walking, all but one (Subject 13 in the original dataset) exhibited lower left push-off force under constraint. Similarly, at 1.1 m/s, 13 participants displayed higher left push-off force during free walking, but only Subject 13 retained this pattern during constrained walking. No participants changed their PO symmetry in the other direction. Note: Fig. 5 does not show this individual data (changes in PO symmetry) for clarity.

The reduction in left-leg propulsion observed in this study mirrors findings in stroke patients, where lower paretic propulsion – analogous to the constrained left leg here – is compensated for by greater non-paretic propulsion, corresponding here to the right leg. With an average PO asymmetry of approximately 80% (Fig. 5, top) and a propulsive impulse asymmetry of 72% (see Appendix), our participants align with stroke patients classified as *mild* to *moderate* in [21]. In their study, mild stroke patients had a propulsive impulse asymmetry of 49%, moderate patients 36%, and severe patients 16%, calculated as the paretic propulsive impulse divided by the total impulse. When converting our symmetry calculation to align with the method in [21], PO asymmetry reported herein corresponds to about 45% and impulse asymmetry to about 42%. The preferred walking speeds of mild to moderate stroke patients in [21] (0.4–1.3 m/s) overlap significantly with the speeds examined here, further reinforcing the relevance of these findings to clinical populations.

Functional asymmetry had the least impact on SL symmetry at the highest walking speed (1.1 m/s) and the most at the lowest speed (0.4 m/s). At 1.1 m/s, the primary change was slightly increased variability in SL symmetry. At 0.4 m/s, SL symmetry shifted on average from slightly longer left steps (symmetry *>*50%) to slightly shorter left steps (symmetry *<*50%), accompanied by a notable increase in inter-person variability (4.5 percentage points). Anecdotally, participants reported feeling *less stable* and walked with the smallest margin of stability at 0.4 m/s [48], potentially explaining these changes. At 0.8 m/s, participants slightly increased SL asymmetry, moving from near-perfect symmetry during free walking to longer left steps during constrained walking, with variability doubling at this speed.

Consistent with findings in patient populations [53, 54], most participants in our cohort walked with equal or longer steps on the constrained (left) leg. However, unlike the slow speeds typically preferred by stroke patients with higher paretic SL [53], participants in our study with SL asymmetry above 52.5% predominantly walked at 0.8 and 1.1 m/s (7/11 subjects; Fig. A3). In other words, increasing speed had a tendency to increase SL asymmetry by taking longer steps with the constrained leg. Interestingly, deviations in SL symmetry, albeit only within a few percentage points of perfect symmetry (50%), mirrored changes in PO symmetry: participants with reduced constrained-leg propulsion (PO symmetry *<*100%) were more likely to take longer left steps. Consequently, the two symmetries correlate (see Fig. A3 in Appendix), similar to the trends identified in patient population by [54].

This suggests a mechanical relationship between step length and propulsive forces, achieved through coordinated control of trunk progression via the stance leg and the timing and positioning of the swing leg. For our participants – most of whom walked with equal or longer left steps despite PO asymmetry – it appears that both mechanisms were engaged. At faster speeds (0.8 and 1.1 m/s), participants took longer steps with their constrained leg. Concurrently, as shown in [48], the higher PO generated by the right leg observed in this study propelled the CoM further forward during the stance phase of the right leg. This forward propulsion strategy is also common in patient populations, enabling them to maintain SL symmetry above 50% (i.e., longer paretic steps) despite propulsive deficits.

Importantly, while propulsive deficits are common in patient populations –regardless of whether they take longer, shorter, or equal steps with the paretic leg – forward CoM progression primarily compensates for this deficit when walking with longer paretic steps. Alternative strategies, such as bilateral hip compensation, are often employed to maintain symmetric steps [53]. While our study did not focus on joint moments or powers, anecdotal reports of hip fatigue among symmetric walkers in our cohort suggest that they may have adopted a similar compensatory mechanism. Further investigation would be needed to confirm this hypothesis.

### 4.4 Cross-dataset predictions of step length and gait propulsion forces

As argued by [10], statistical gait models can serve as individualised biofeedback targets for patients, offering insights into gait pattern deficits and their magnitudes. Unlike machine learning models, statistical models provide transparency by revealing not only if, but *how*, independent gait variables influence the dependent one (model output). However, model performance reflects both the model design and the underlying gait dynamics; consequently, multiple cross-dataset predictions are necessary to disentangle the two factors.

Models developed by [10] were trained on the same F18 dataset as used herein^3^. Their model evaluations done using Aikake Information Criterion (AIC) showed that gait speed alone is sufficient to predict peak push-off force (PO). Our findings validate this for both F18 and B24 datasets, as shown in Table A2. Importantly, we demonstrate in the same table how including different gait parameters and participant cohorts has a huge impact on model estimation quality. For example, constrained walking data from B24 and the Elderly cohort from F18 posed significant challenges for model estimation, while models trained on B24 free walking data performed best. This improvement was especially pronounced when accounting for multiple cadences per speed (*Model1 ↓Model2*), highlighting the utility of cadence as a model input variable.

Cross-dataset predictions of PO presented herein (B24*↓*F18, and vice versa; Fig. A5) deliver slightly worse performance as measured by *R*^2^ compared to the result presented in [10] (*R*^2^ = 0.76). However, several important points must be considered. First, our models were trained on a narrower range of speeds and evaluated on fewer data points, which likely reduced *R*^2^. Second, our cross-dataset predictions handled F18 Young and Elderly separately, and these two datasets present different modelling challenges (see Table A2). Third, when we trained models on and predicted data using the full F18 dataset – similar to [10], i.e., combining Young and Elderly cohorts and using the full range of speeds – the results were comparable: F18 predicting B24 achieved an *R*^2^ of 0.70 using speed alone (*Model1*) and 0.79 using speed and cadence (*Model2*) (but 0.68 using *Model3*). However, the opposite direction – predicting full F18 from B24 – showed a significant drop in accuracy, with *R*^2^ values of 0.28 for *Model1* and 0.38 for *Model2* (and 0.39 for *Model3*; note that these predictions are not visualised in this paper). When we predicted F18 within the range of speeds present in B24 but still combined Young and Elderly cohorts, the *R*^2^ only slightly increased (to 0.4 for *Model1* and 0.38 for *Model2* and *Model3*), suggesting that while the range of tested speeds is important, the differences in population cohorts have a more pronounced impact on prediction accuracy.

Similar to PO models, SL model estimation performed well using speed alone as a model input, with AIC evaluations confirming that anthropometric variables add minimal value to model estimation quality (Table A4). The inclusion of trailing limb angle (TA) as an additional variable (*Model2*) substantially improved SL model estimation in the B24 dataset, where multiple cadences per speed were tested – an improvement analogous to the role of cadence in PO modelling. These findings align with those in [10], which also demonstrated the sufficiency of gait speed as a primary predictor for SL and highlighted challenges in modeling data from the Elderly cohort. Unlike PO estimation, constraints had a less pronounced impact on SL estimation quality. Remarkably, the B24

*Constrained Right* dataset (*Model2* and *Model3*) yielded the highest fidelity SL models across all conditions, significantly outperforming models trained on F18 data. This suggests that SL dynamics may be less sensitive to constraints compared to PO, possibly due to compensatory mechanisms that stabilise spatial gait parameters even under altered gait conditions.

SL predictions within the F18 dataset, from Young to Elderly and vice versa, showed comparable performance to the cross-dataset predictions reported by [10], with *R*^2^ values around 0.7 (*Model1*, Fig. A6). Cross-dataset predictions from B24 to F18 using *Model1* performed slightly worse when predicting the Elderly cohort (*R*^2^ = 0.66) but achieved much better accuracy when predicting the Young cohort (*R*^2^ = 0.86), as shown in Fig. A7. Predicting the combined Young and Elderly cohorts from a B24-trained *Model1* yielded intermediate accuracy (*R*^2^ = 0.77), closely aligning with SL prediction performance in [10] when the speed range was consistent between training and testing datasets. Notably, the significant discrepancy in SL prediction accuracy between the Young and Elderly cohorts, which is not observed in PO predictions, prompted additional analyses incorporating age as a variable.

As demonstrated in Fig. A8, age did not improve predictions between B24 and F18 Young cohorts (or vice versa), as expected due to the similar age ranges of these groups. Surprisingly, adding age to the model worsened B24 *↓* F18 Elderly predictions, a pattern consistent across multiple predictions tested in this study, where adding anthropometric variables often reduced cross-dataset prediction accuracy (these results are not visualised due to the lack of space). However, adding age significantly improved predictions from F18 Elderly to B24, suggesting that age plays a more critical role in models trained on elderly cohorts, likely capturing age-related changes in SL dynamics.

Overall, these results demonstrate that while statistical models can reasonably predict SL and PO across datasets, their performance is highly dependent on the range and characteristics of the training data. Variations in population dynamics (e.g., age, cadence variability) and experimental design (e.g., speed ranges, constraints) highlight the need for diverse and representative datasets to improve model generalisability. Importantly, the findings also caution against over-reliance on anthropometric variables, as these can enhance estimation metrics like AIC but frequently reduce cross-dataset prediction performance, underscoring the need for rigorous model evaluation.

### 4.5 Limitations

This study provides invaluable insights into human gait adaptations, but it does not come without limitations. First, our cohort comprises only healthy young adults, limiting the generalisability of the presented findings. Although our experimental design intentionally excludes confounding factors common in patient populations, such as reduced strength or neuromuscular control, it is possible that elderly participants or individuals with other impairments would respond differently to imposed functional asymmetry. Second, the passive knee brace used to induce asymmetry and emulate hemiparetic gait does not replicate the biomechanical complexities of pathological conditions, such as stroke, where spasticity and sensory deficits play a significant role in gait adaptations. Third, while the study examines a wide range of walking speeds and cadences, it excludes extreme speeds typically found in clinical populations (e.g., 0.2 m/s and 1.3 m/s), where distinct gait dynamics are known to govern gait patterns. Finally, our focus on short-term gait adaptations within five minutes of each condition leaves longer-term compensatory strategies and motor learning unexplored and unknown. Addressing these limitations in future research would enhance the clinical relevance and broader applicability of these findings.

## 5 Conclusion

This study explored how healthy young adults adapt their gait in response to functional asymmetry induced by a unilateral knee constraint, focusing on step length (SL) and push-off force (PO) across varying walking speeds and cadences. The findings reveal distinct adaptation strategies, with participants prioritising spatial over kinetic symmetry, particularly at higher speeds. While SL remained relatively stable across conditions, propulsive force on the constrained leg decreased significantly, prompting compensatory increases in the non-constrained leg’s propulsion. These adaptations align with patterns observed in hemiparetic populations, suggesting shared biomechanical strategies for managing asymmetry.

The results underscore the importance of understanding individual gait adaptations to functional asymmetry, offering valuable insights for rehabilitation strategies in clinical populations. By isolating the effects of a constrained joint on walking dynamics, this study provides a framework for future investigations into the mechanisms underlying asymmetrical gait and highlights the role of biomechanical trade-offs in maintaining efficiency and stability during walking. Further research integrating joint moment analysis and overground walking conditions is necessary to refine these insights and broaden their application to diverse patient populations.

## Author Contributions

T.B. ideated, prepared, and led experimental study and this work, extracted and pre-processed raw data, analysed data in Matlab, drafted all manuscript versions, prepared (most of the) figures, tables, and data visualisation, and secured funding (ECRG143) to support data analysis in this work. Y.X. carried out statistical data analysis and helped with preparing figures, tables, and data visualisation. L.P. contributed his expertise to experimental study design and guided statistical modelling and data analysis. D.O. and Y.T. provided feedback on the manuscript and contributed to developing methodology, data analysis and interpretation, project administration and supervision, and secured funding for the data collection project.

## Acknowledgments

The authors would like to thank Prof. Gavin Williams, PhD FACP, from Epworth Hospital and The University of Melbourne, for his support in the study’s preparation and his expertise in interpreting the results and situating them within a broader context.

## Data Availability Statement

The two datasets used in this study can be found in figshare repositories: Baček et al. dataset at [55], and Fukuchi et al. dataset at [56].

## Footnotes

1 R-Squared: represents the proportion of variance in the dependent variable predictable from the independent variables. MAE: provides the average magnitude of prediction errors, disregarding direction. MAPE: measures accuracy as a percentage, offering an intuitive view of model performance.

2 Temporal gait aspects, including step time, swing time, and double stance time, as well as their symmetries will be analysed in our future work.

3 Note that [10] used the full range of speeds and combined Young and Elderly data in all their models

## References

[1] H. J. Ralston, “Energy-speed relation and optimal speed during level walking.,” Int. Z. Angew. Physiol. Einschl. Arbeitsphysiol., vol. 17, pp. 277–283, 4 1958, issn: 0020-9376 (Print).

[2] N. H. Molen, R. H. Rozendal, and W Boon, “Graphic representation of the relationship between oxygen-consumption and characteristics of normal gait of the human male.,” Proc K Ned Akad Wet C., vol. 75, pp. 305–314, 4 1972.

[3] J. M. Donelan, R Kram, and A. D. Kuo, “Mechanical and metabolic determinants of the preferred step width in human walking.,” Proc. R. Soc. B Biol. Sci., vol. 268, pp. 1985–1992, 1480 Oct. 2001, issn: 0962-8452.

[4] A. Minetti, C Capelli, P Zamparo, P. Prampero, and F Saibene, “Effects of stride frequency on mechanical power and energy expenditure of walking,” Medicine and science in sports and exercise, vol. 27, pp. 1194–1202, 8 1995, issn: 0195-9131.

[5] R. Ellis, K. Howard, and R. Kram, “The metabolic and mechanical costs of step time asymmetry in walking,” Proceedings. Biological sciences / The Royal Society, vol. 280, p. 20 122 784, Apr. 2013.

[6] H. Reimann, T. Fettrow, and J. J. Jeka, “Strategies for the control of balance during locomotion,” Kinesiology Review, vol. 7, pp. 18–25, 1 2018.

[7] M. Orendurff, A. D. Segal, G. K. Klute, J. S. Berge, E. Rohr, and N. J. Kadel, “The effect of walking speed on center of mass displacement.,” Journal of rehabilitation research and development, vol. 41 6A, pp. 829–34, 2004.

[8] S. Bruijn and J. V. Dieen, “Control of human gait stability through foot placement,” Journal of The Royal Society Interface, vol. 15, p. 20 170 816, Jun. 2018.

[9] J. M. Donelan, R. Kram, and A. D. Kuo, “Mechanical work for step-to-step transitions is a major determinant of the metabolic cost of human walking,” J. Exp. Biol., vol. 205, pp. 3717– 3727, 23 2002, issn: 0022-0949.

[10] M. B. Yanez, S. A. Kettlety, J. M. Finley, N. Schweighofer, and K. A. Leech, “Gait speed and individual characteristics are related to specific gait metrics in neurotypical adults,” Scientific Reports, vol. 13, p. 8069, 1 2023, issn: 2045-2322.

[11] L. Hak, H. Houdijk, P. Beek, and J. V. Dieen, “Steps to take to enhance gait stability: The effect of stride frequency, stride length, and walking speed on local dynamic stability and margins of stability,” PloS one, vol. 8, e82842, Dec. 2013.

[12] J. R. Franz and R. Kram, “The effects of grade and speed on leg muscle activations during walking,” Gait Posture, vol. 35, pp. 143–147, 1 2012, issn: 0966-6362.

[13] H. Xie and J. H. Chien, “Walking on different inclines affects gait symmetry differently in the anterior-posterior and vertical directions: Implication for future sensorimotor training,” PeerJ, vol. 12, R. Baptista, Ed., e18096, 2024, issn: 2167-8359.

[14] H. B. Frame, C. Finetto, J. C. Dean, and R. R. Neptune, “The influence of lateral stabilization on walking performance and balance control in neurologically-intact and post-stroke individuals,” Clinical Biomechanics, vol. 73, pp. 172–180, 2020, issn: 0268-0033.

[15] T. M. Owings and M. D. Grabiner, “Step width variability, but not step length variability or step time variability, discriminates gait of healthy young and older adults during treadmill locomotion,” Journal of Biomechanics, vol. 37, pp. 935–938, 6 2004, issn: 0021-9290.

[16] J. C. Dean, N. B. Alexander, and A. D. Kuo, “The effect of lateral stabilization on walking in young and old adults,” IEEE Transactions on Biomedical Engineering, vol. 54, pp. 1919–1926, 11 2007, issn: 1558-2531.

[17] M Brandstater, H. Debruin, C Gowland, and B Clark, “Hemiplegic gait: Analysis of temporal variables,” Archives of physical medicine and rehabilitation, vol. 64, pp. 583–587, Jan. 1984.

[18] S. J. Olney and C. Richards, “Hemiparetic gait following stroke. part i: Characteristics,” Gait Posture, vol. 4, pp. 136–148, 2 1996, issn: 0966-6362.

[19] G Stoquart, C Detrembleur, and T. M. Lejeune, “The reasons why stroke patients expend so much energy to walk slowly.,” Gait Posture, vol. 36, pp. 409–413, 3 Jul. 2012, issn: 1879-2219 (Electronic).

[20] C. Kim and J. J. Eng, “Symmetry in vertical ground reaction force is accompanied by symmetry in temporal but not distance variables of gait in persons with stroke,” Gait Posture, vol. 18, pp. 23–28, 1 2003, issn: 0966-6362.

[21] M. G. Bowden, C. K. Balasubramanian, R. R. Neptune, and S. A. Kautz, “Anterior-posterior ground reaction forces as a measure of paretic leg contribution in hemiparetic walking,” Stroke, vol. 37, pp. 872–876, 3 Mar. 2006.

[22] J. C. Wall and G. I. Turnbull, “Gait asymmetries in residual hemiplegia.,” Archives of physical medicine and rehabilitation, vol. 67 8, pp. 550–3, 1986.

[23] A.-L. Hsu, P.-F. Tang, and M.-H. Jan, “Analysis of impairments influencing gait velocity and asymmetry of hemiplegic patients after mild to moderate stroke,” Archives of Physical Medicine and Rehabilitation, vol. 84, pp. 1185–1193, 8 Aug. 2003, issn: 0003-9993.

[24] K. K. Patterson, W. H. Gage, D. Brooks, S. E. Black, and W. E. McIlroy, “Evaluation of gait symmetry after stroke: A comparison of current methods and recommendations for standardization,” Gait Posture, vol. 31, pp. 241–246, 2 2010, issn: 0966-6362.

[25] M. D. Lewek, C. Raiti, and A. Doty, “The presence of a paretic propulsion reserve during gait in individuals following stroke,” Neurorehabilitation and Neural Repair, vol. 32, pp. 1011–1019, 12 Dec. 2018, issn: 1545-9683.

[26] P. Padmanabhan, K. S. Rao, S. Gulhar, K. M. Cherry-Allen, K. A. Leech, and R. T. Roemmich, “Persons post-stroke improve step length symmetry by walking asymmetrically,” Journal of NeuroEngineering and Rehabilitation, vol. 17, p. 105, 1 2020, issn: 1743-0003.

[27] L. Hak et al., “Stepping strategies used by post-stroke individuals to maintain margins of stability during walking,” Clinical Biomechanics, vol. 28, pp. 1041–1048, 9 2013, issn: 0268-0033.

[28] M. D. Lewek, C. E. Bradley, C. J. Wutzke, and S. M. Zinder, “The relationship between spatiotemporal gait asymmetry and balance in individuals with chronic stroke,” Journal of Applied Biomechanics, vol. 30, pp. 31–36, 1 2014.

[29] Y. P. Lim, Y.-C. Lin, and M. G. Pandy, “Lower-limb muscle function in healthy young and older adults across a range of walking speeds.,” Gait Posture, vol. 94, pp. 124–130, 2022, issn: 0966-6362.

[30] M. J. Booij, E Meinders, I. N. Sierevelt, P. A. Nolte, J Harlaar, and J. C. van den Noort, “Matching walking speed of controls affects identification of gait deviations in patients with a total knee replacement.,” Clinical Biomechanics, vol. 82, Feb. 2021, issn: 0268-0033.

[31] S. A. Roelker, M. G. Bowden, S. A. Kautz, and R. R. Neptune, “Paretic propulsion as a measure of walking performance and functional motor recovery post-stroke: A review,” Gait Posture, vol. 68, pp. 6–14, 2019, issn: 0966-6362.

[32] T. Baček et al., “A biomechanics and energetics dataset of neurotypical adults walking with and without kinematic constraints,” Scientific Data, vol. 11, p. 646, 1 2024, issn: 2052-4463.

[33] D. Winter, Biomechanics and Motor Control of Human Movement. 4th. Wiley, Sep. 2009.

[34] M. J. Lindstrom and D. M. Bates, “Newton-raphson and em algorithms for linear mixed-effects models for repeated-measures data,” Journal of the American Statistical Association, vol. 83, pp. 1014–1022, 404 1988.

[35] C. A. Fukuchi, R. K. Fukuchi, and M. Duarte, “Effects of walking speed on gait biomechanics in healthy participants: A systematic review and meta-analysis.,” Systematic reviews, vol. 8, p. 153, 1 Jun. 2019, issn: 2046-4053 (Electronic).

[36] C. A. Fukuchi, R. K. Fukuchi, and M. Duarte, “A public dataset of overground and treadmill walking kinematics and kinetics in healthy individuals,” PeerJ, vol. 6, M. Zago, Ed., 2018, issn: 2167-8359. doi: 10.7717/peerj.4640.

[37] D. Robertson, G. E. Caldwell, J. Hamill, G. Kamen, and S. N. Whittlesey, Research Methods in Biomechanics: Second edition (eBook). Nov. 2013, isbn: 0-7360-8340-0.

[38] G. Wu et al., “Isb recommendation on definitions of joint coordinate system of various joints for the reporting of human joint motion—part i: Ankle, hip, and spine,” Journal of Biomechanics, vol. 35, pp. 543–548, 4 2002, issn: 0021-9290.

[39] S. J. Ali, A. N. Ansari, A Rahman, S Imtiyaz, and B Rashid, “Post-stroke hemiplegic gait: A review,” Pharma Innov., vol. 3, pp. 36–41, 8 2014.

[40] T. Baček et al., “Varying joint patterns and compensatory strategies can lead to the same functional gait outcomes: A case study,” in 2022 International Conference on Rehabilitation Robotics (ICORR), 2022, pp. 1–6.

[41] B. F. Mentiplay, M. Banky, R. A. Clark, M. B. Kahn, and G. Williams, “Lower limb angular velocity during walking at various speeds,” Gait Posture, vol. 65, pp. 190–196, 2018, issn: 0966-6362.

[42] S. Park, C. Liu, N. Sánchez, J. K. Tilson, S. J. Mulroy, and J. M. Finley, “Using biofeedback to reduce step length asymmetry impairs dynamic balance in people poststroke,” Neurorehabilitation and Neural Repair, vol. 35, pp. 738–749, 8 Jun. 2021, issn: 1545-9683.

[43] J. L. Lelas, G. J. Merriman, P. O. Riley, and D. Kerrigan, “Predicting peak kinematic and kinetic parameters from gait speed,” Gait Posture, vol. 17, pp. 106–112, 2 2003, issn: 0966-6362.

[44] C. A. Fukuchi, R. K. Fukuchi, and M. Duarte, “Test of two prediction methods for minimum and maximum values of gait kinematics and kinetics data over a range of speeds,” Gait Posture, vol. 73, pp. 269–272, 2019, issn: 0966-6362.

[45] C. A. Fukuchi and M. Duarte, “A prediction method of speed-dependent walking patterns for healthy individuals,” Gait Posture, vol. 68, pp. 280–284, 2019, issn: 0966-6362.

[46] E. Y. Chao, R. K. Laughman, E Schneider, and R. N. Stauffer, “Normative data of knee joint motion and ground reaction forces in adult level walking,” Journal of Biomechanics, vol. 16, pp. 219–233, 3 1983, issn: 0021-9290.

[47] T. Oberg, A. Karsznia, and K. Oberg, “Basic gait parameters: Reference data for normal subjects, 10–79 years of age,” Journal of rehabilitation research and development, vol. 30, pp. 210–223, 1993.

[48] T. Baček, D. Oetomo, and Y. Tan, “Gait adaptations under functional asymmetry: Exploring the role of step width, step length, and com in lateral stability,” bioRxiv, 2024. [Online]. Available: https://www.biorxiv.org/content/early/2024/12/23/2024.12.23.630028.

[49] A. D. Kuo, “A simple model of bipedal walking predicts the preferred speed-step length relationship,” J. Biomech. Eng., vol. 123, pp. 264–269, 3 Jan. 2001, issn: 0148-0731.

[50] J. D. Polk, R. M. Stumpf, and K. S. Rosengren, “Limb dominance, foot orientation and functional asymmetry during walking gait,” Gait Posture, vol. 52, pp. 140–146, 2017, issn: 0966-6362.

[51] M. K. Seeley, B. R. Umberger, and R. Shapiro, “A test of the functional asymmetry hypothesis in walking,” Gait Posture, vol. 28, pp. 24–28, 1 2008, issn: 0966-6362.

[52] J. Stenum and J. T. Choi, “Disentangling the energetic costs of step time asymmetry and step length asymmetry in human walking,” Journal of Experimental Biology, vol. 224, 12 Jun. 2021, issn: 0022-0949.

[53] J. L. Allen, S. A. Kautz, and R. R. Neptune, “Step length asymmetry is representative of compensatory mechanisms used in post-stroke hemiparetic walking.,” Gait posture, vol. 33, pp. 538–543, 4 Apr. 2011, issn: 1879-2219 (Electronic).

[54] C. K. Balasubramanian, M. G. Bowden, R. R. Neptune, and S. A. Kautz, “Relationship between step length asymmetry and walking performance in subjects with chronic hemiparesis,” Archives of Physical Medicine and Rehabilitation, vol. 88, pp. 43–49, 1 Jan. 2007, issn: 0003-9993.

[55] T. Baček et al., “A biomechanics and energetics dataset of healthy young adults walking with and without kinematic constraints.,” figshare, 2023. doi: https://figshare.com/s/b625dafe53f4f83e21cd.

[56] C. A. Fukuchi, R. K. Fukuchi, and M. Duarte, “A public dataset of overground and treadmill walking kinematics and kinetics in healthy individuals.,” PeerJ, vol. 6, M. Zago, Ed., e4640, 2018, issn: 2167-8359. doi: 10.6084/m9.figshare.5722711.v2.

