## Supplementary file for "Gait Adaptations in Step Length and Push-off Force during Walking with Functional Asymmetry"

### Supplemental Data

Here, we provide more figures pertaining to vertical (weight bearing) and fore-aft (propulsive) ground reaction force (GRF) impulse, peak push-off force and step length asymmetry correlation, and evaluate bi-directional cross-dataset model predictions, using B24 and [36] (hereafter called F18) datasets.

#### Vertical and propulsive impulse

The vertical ground reaction force (GRF) impulse for each leg across all walking conditions is shown in Fig. A1, while the fore-aft (propulsive) GRF impulse is shown in Fig. A2. During free walking (blue bars), there is a slight tendency toward higher vertical impulses on the right leg, as reflected in the vertical GRF symmetry (calculated as the ratio of left to right impulses, with 100% = symmetric): the average across five cadences per speed is 97.2±0.6% at 1.1 m/s; 97.9±0.2% at 0.8 m/s; and 97.8±0.1% at 0.4 m/s. In contrast, fore-aft (propulsive) impulses tend to be slightly higher on the left leg, as indicated by the propulsive GRF symmetry: the average across five cadences per speed is 102.45±0.74% at 1.1 m/s, 101.15±2.9% at 0.8 m/s, and 104.1±2.9% at 0.4 m/s.

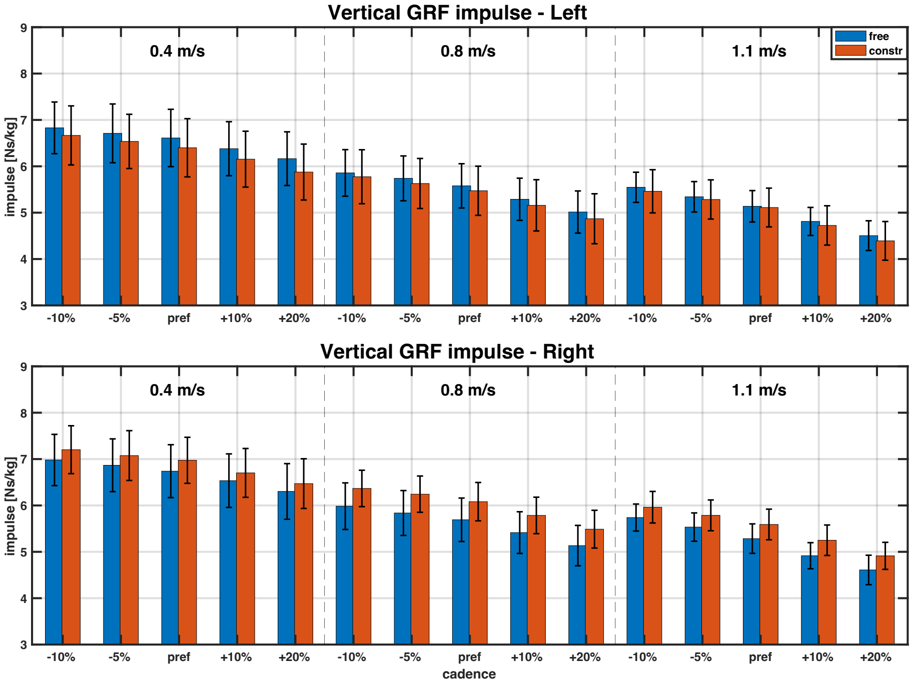


Figure A1: Vertical (weight bearing) ground reaction force (GRF) impulse. Data are average across all participants. Three walking speeds are separated by vertical dashed lines, and two conditions (free, constrained) are colour-coded. Note same y-axis limits on both graphs.

During constrained walking (red bars), the vertical GRF impulse decreased on the left leg and increased on the right leg compared to free walking. This resulted in a further increase in vertical

GRF impulse asymmetry, shifting more toward the right side. The average symmetry ratio across five cadences per speed was 90.7±1.0% at 1.1 m/s; 89.6±0.8% at 0.8 m/s; and 91.8±0.7% at 0.4 m/s. The asymmetry shift was much more pronounced in the fore-aft GRF impulse particularly at higher speeds. At 0.4 m/s, fore-aft GRF impulse asymmetry remained similar to free walking, with average ratio of 102.3±8.1% across all cadences. However, at higher speeds, the asymmetry direction reversed during constrained walking. The average ratio across five cadences per speed was 72.9±3.4% at 1.1 m/s and 71.55±4.1% at 0.8 m/s, indicating a marked reduction in left-leg propulsion relative to the right leg.

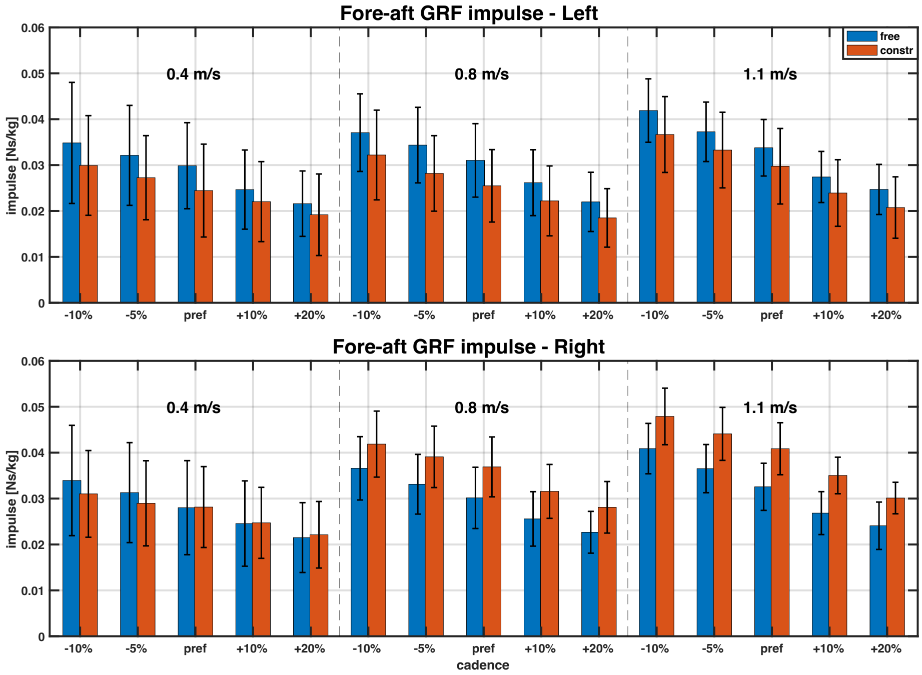


Figure A2: Fore-aft (propulsive) ground reaction force (GRF) impulse. Data are average across all participants. Three walking speeds are separated by vertical dashed lines, and two conditions (free, constrained) are colour-coded. Note same y-axis limits on both graphs.

#### Peak push-off force and step length symmetry correlation

Fig. A3 illustrates the relationship between changes in PO symmetry and SL symmetry during free and constrained walking. To analyse this correlation, we used ordinary least squares (OLS) regression, treating SL symmetry as the independent (exogenous) variable and PO symmetry as the dependent (endogenous) variable. The analysis was conducted separately for free and constrained walking, with correlations tested both at each individual speed (19 data points, one per participant) and across all three speeds combined (57 data points: 3 speeds × 19 participants).

In all cases, there was a significant correlation between the two symmetries. During free walking, the adjusted $R^2$ was higher than 0.98 across all four conditions, with the model parameter (no intercept) estimated at 0.0207±7.3e$^{-4}$. During constrained walking, the adjusted $R^2$ was slightly lower compared to free walking, but remained above 0.94 across all four conditions. The model parameter (no intercept) was estimated at 0.0169±0.002.

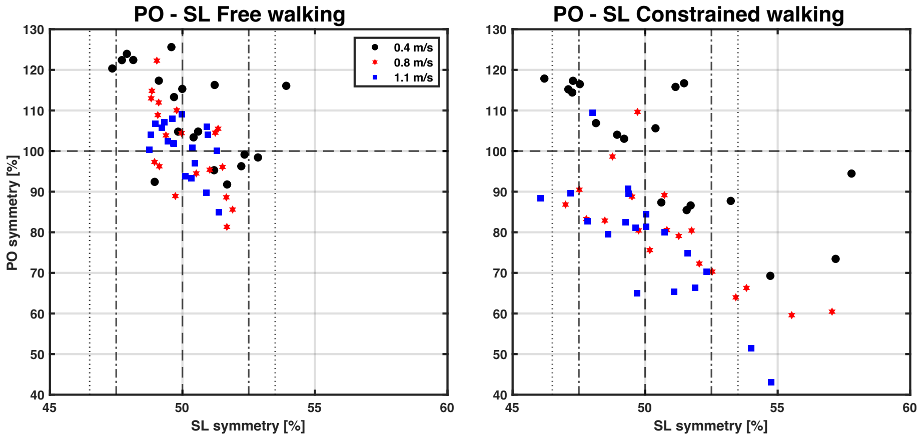


Figure A3: Relationship between peak push-off force (PO) symmetry and step length (SL) symmetry. Left subfigure shows PO-SL relationship during free walking, and right subfigure during constrained walking. Each datapoint is one person per speed (i.e., average value over 5 step frequencies). Different colours and marker shapes are used to separate walking speed. Horizontal dashed line (100%) corresponds to perfect PO symmetry, and vertical dashed line (50%) to perfect SL symmetry. Dosh-dotted vertical lines (47.5% and 52.5%) correspond to ≈10% difference in step length for direct comparison with [54] (Fig. 3 in their paper; step length ratio – SLR – of 0.9 and 1.1, respectively), and dotted vertical lines (46.5% and 53.5%) correspond to SL symmetry definition from [53].

#### Fukuchi et al. dataset (F18)

A 2018 dataset by Fukuchi et al. includes data from 42 neurotypical adults, comprising 24 young adults (age 28±4 years, body mass 68.4±12.2 kg, height 171.1±10.5 cm) and 18 older adults (age 63±8 years, body mass 66.9±10.1 kg, and height 161.8±9.5 cm). All participants were free of any lower-extremity injury and walked barefoot. Trials took place on an instrumented treadmill, each lasting 90 seconds, with the final 30 seconds used for analysis. Each participant completed eight walking trials, ranging from 40% to 145% of their self-selected speed in 15% increments, presented in a randomised order. The mean self-selected speed was 1.25±0.16 m/s, with a comfortable gait speed ranging from 0.89 to 1.54 m/s, and an overall range of speeds from 0.36 to 2.23 m/s. Leg length was provided with the dataset, measured as the distance between the anterior superior iliac spine (ASIS) and the ipsilateral medial malleolus in a supine position. Kinematic data were recorded at 150 Hz and kinetic data at 300 Hz.

Participants were allowed to use handrails during walking, which could affect kinetic data. For this reason, we excluded four elderly participants (numbers 27, 32, 37, and 39 in the original dataset) who used handrails. Additionally, one participant (number 33 in the original dataset) was removed due to major offsets in their force data, and three more participants (numbers 40, 41, and 42 in the original dataset) were removed for exhibiting unrealistic vertical force values. Consequently, we trained our models with all 24 young adults (*F18 Young*) and a reduced sample of 10 older participants (*F18 Elderly*), resulting in the anthropometric features detailed in Table A1. For direct comparison with the Bacek et al. dataset, we only used the first four speeds from the Fukuchi et al. dataset. In cases where full eight-speed datased was used, we denote this with an asterisk (e.g., *F18 Young**).

Some trials exhibited overstepping (e.g., left leg stepping onto the right belt), and these trials were excluded from the analysis. Specifically, we excluded Test 08 (fastest speed) for participants 28, 29, 36, and 38; Test 02 and 03 (second and third speeds) for participant 35; and Test 07 for participant 38. The remaining trials from these participants were retained since our statistical approach, described in the main paper, is robust to missing data. Given the high correlation between gait parameters in the left and right legs, only left leg data from the Fukuchi et al. dataset was used for analysis.

Table A1: Anthropometric characteristics of participants in two datasets used in this work.

| | Fukuchi et al. 2018 | | Bacek et al. 2024 |
|---|---|---|---|
| | Young | Elderly | Young |
| **# of participants** | 24 | 10 | 19 |
| **age** *[years]* | 26 ± 4 | 61 ± 6 | 30 ± 8 |
| **sex** *[Male/Female]* | 10/24 female | 4/10 female | 5/19 female |
| **body mass** *[kg]* | 68.4 ± 12.2 | 65.4 ± 10.2 | 72.6 ± 12.6 |
| **leg length** *[cm]* | 87.8 ± 5.9 | 87 ± 6.9 | 88.2 ± 6.3 |
| **speed range** *[m/s]* | 0.39 - 1.31 | 0.42 - 1.24 | 0.4 - 1.1 |
| **speed range*** *[m/s]* | 0.39 - 2.23 | 0.42 - 1.98 | / |

*Fukuchi et al. dataset with * contains all eight speeds; the one without *, only the first four*

#### Model estimation (within-dataset) quality

##### Peak push-off force

Table A2 presents the model estimation quality for peak push-off force (PO) as assessed by Aikake Information Criterion (AIC). AIC values are provided for both the full F18 dataset (denoted with *), and the F18 subset spanning similar range of walking speeds as the B24 dataset. As discussed in the main paper, PO in B24 is well modelled using only walking speed and cadence (*Model2*); this is validated in Table A3, which shows that only speed and cadence are relevant across all four groups of data. Cadence is expected to be significant in modelling B24 dataset since, as the experimental design includes five cadences per speed, resulting in varied peak push-off force values per speed. Note that the AIC values for B24 in Table A2 are identical to those in the main paper (Table 1), but are repeated here for direct comparison.

PO modelling in F18 Young closely resembles that of the B24 dataset (Table A2) – adding cadence to *Model1* improves estimation accuracy. However, model quality in F18 Young never reaches that

Table A2: Peak push-off force (PO) model evaluation using AIC.

| | Data group | Model1 | Model2 | Model3 |
|---|---|---|---|---|
| B24 | Free Left | -162.18 | -260.44 | -270.64 |
| | Free Right | -192.03 | -270.77 | -283.26 |
| | Constrained Left | -13.265 | -94.35 | -94.11 |
| | Constrained Right | -93.86 | -175.76 | -182.54 |
| F18 | Young | -162.82 | -175.44 | -182.67 |
| | Elderly | -43.04 | -41.56 | -44.82 |
| F18* *(full)* | Young | -162.62 | -181.88 | -197.95 |
| | Elderly | -73.20 | -73.47 | -85.32 |

*Model1 = fcn(v,v²); Model2 = fcn(v,v²,f); Model3 = fcn(v,v²,f,anthropometry)*

Table A3: Peak push-off force (PO) model parameter significance in *Model3*.

| | B24 | | | | F18 | | F18* | |
|---|---|---|---|---|---|---|---|---|
| | Free L | Free R | Const. L | Constr. R | Young | Elderly | Young | Elderly |
| intercept | 0.17 | 0.046 | 0.000 | 0.868 | 0.646 | 0.409 | 0.105 | 0.934 |
| speed (v) | 0.000 | 0.000 | 0.006 | 0.000 | 0.000 | 0.001 | 0.000 | 0.000 |
| speed (v²) | 0.002 | 0.014 | 0.002 | 0.415 | 0.479 | 0.930 | 0.000 | 0.000 |
| cadence | 0.000 | 0.000 | 0.000 | 0.000 | 0.000 | 0.196 | 0.000 | 0.161 |
| leg length | 0.758 | 0.439 | 0.023 | 0.121 | 0.285 | 0.004 | 0.003 | 0.000 |
| sex | 0.006 | 0.523 | 0.240 | 0.087 | 0.014 | 0.845 | 0.059 | 0.513 |
| age | 0.286 | 0.044 | 0.249 | 0.702 | 0.125 | 0.028 | 0.165 | 0.000 |
| weight | 0.017 | 0.046 | 0.000 | 0.868 | 0.646 | 0.409 | 0.105 | 0.934 |

*L = Left; R = Right; F18 in 0.36-1.31 m/s range; F18* in 0.36-2.23 m/s range; p<0.05 = significant (in green)*

of the best B24 model (*Free Right*), even when anthropometric variables are included (*Model3*). It is worth noting that participants in the B24 and F18 Young datasets have similar anthropometric characteristics. Conversely, PO modelling for the F18 Elderly data shows no improvement with the addition of cadence, and adding anthropometric variables has minimal impact, with Age as the only exception. This is confirmed in Table A3, where the F18 Elderly model is the only one with insignificant cadence parameter and a significant Age parameter. Interestingly, when modelling the full range of speeds in F18 (Young* and Elderly*), $v^2$ becomes significant, unlike when modelling a reduced range of speeds (F18 Young and Elderly). In the B24 dataset, $v^2$ is relevant even in a smaller range of speeds (0.4 - 1.1 m/s).

#### Step length

Table A4 shows model estimation quality for step length (SL) assessed by AIC. Similar to PO, AIC values are provided for both the full F18 dataset (denoted with *) and the F18 subset that spans a similar range of speeds as the B24 dataset. (Note: AIC values are not available for F18 *Model2* since trailing limb angle is not included in F18 due to its unavailability with the dataset; also, *Model3* includes trailing limb for B24 but not F18). As discussed in the main paper, SL in B24 is effectively modelled using only walking speed and trailing limb angle (*Model2*). This is confirmed in Table A5, showing that only these parameters significantly contribute to model estimation quality across all B24 groups. Note that F18 *Model3* in Table A4 only includes walking speed and anthropometry, while B24 also includes trailing limb angle. The AIC values for B24 in Table A4 are the same as in the main paper (Table 1), but are repeated here for direct comparison.

Table A4: Step length (SL) model evaluation using AIC.

| | Data group | Model1 | Model2 | Model3 |
|---|---|---|---|---|
| B24 | Free Left | -782.61 | -1008.58 | -1036.79 |
| | Free Right | -752.03 | -958.57 | -985.08 |
| | Constrained Left | -709.44 | -1036.96 | -1055.63 |
| | Constrained Right | -746.34 | -853.99 | -870.60 |
| F18 | Young | -435.10 | NA | -432.32 |
| | Elderly | -157.56 | NA | -152.62 |
| F18* *(full)* | Young | -853.32 | NA | -849.17 |
| | Elderly | -317.05 | NA | -314.09 |

*Model1 = fcn($v$,$v^2$); Model2 = fcn($v$,$v^2$,TA);, Model3 = fcn($v$,$v^2$,TA,anthropometry)*

Table A5: Step length (SL) model parameter significance in *Model3*.

| | B24 | | | | F18 | | F18* | |
|---|---|---|---|---|---|---|---|---|
| | Free L | Free R | Const. L | Constr. R | Young | Elderly | Young | Elderly |
| intercept | 0.000 | 0.000 | 0.078 | 0.275 | 0.239 | 0.022 | 0.013 | 0.001 |
| speed (v) | 0.000 | 0.000 | 0.000 | 0.000 | 0.000 | 0.000 | 0.000 | 0.000 |
| speed ($v^2$) | 0.005 | 0.013 | 0.000 | 0.007 | 0.083 | 0.799 | 0.000 | 0.000 |
| leg length | 0.000 | 0.000 | 0.170 | 0.763 | 0.217 | 0.883 | 0.733 | 0.192 |
| sex | 0.000 | 0.000 | 0.056 | 0.02 | 0.972 | 0.888 | 0.980 | 0.356 |
| age | 0.016 | 0.116 | 0.001 | 0.308 | 0.494 | 0.197 | 0.102 | 0.398 |
| trail angle | 0.000 | 0.000 | 0.000 | 0.000 | / | / | / | / |
| weight | 0.512 | 0.115 | 0.463 | 0.002 | 0.269 | 0.368 | 0.547 | 0.509 |

*L = Left; R = Right; F18 in 0.36-1.31 m/s range; F18* in 0.36-2.23 m/s range*

SL modelling in F18 closely resembles that in B24 for both for Young and Elderly groups (in contrast to PO, which aligned only with F18 Young). Adding anthropometric variables does not significantly enhance model estimation quality, and once again, the best quality model within the 0.4-1.1 m/s range of speeds is from the B24 dataset (*Constrained Left*). Interestingly, even when considering a broader speed range in F18 Young (*F18**), F18 *Model1* only slightly outperforms best B24 model (*Free Left*), despite using a range of speeds twice as large. Notably, when the range of speed is doubled in F18 dataset (i.e., *F18**), both Young and Elderly models start to rely heavily on $v^2$ in their modelling (see Table A5). Within the narrower speed range of the B24 dataset, $v^2$ is significant for F18 Young and B24 models, while F18 Elderly does not rely on $v^2$ at all.

#### Bi-directional cross-dataset predictions

##### Peak push-off force

We begin by establishing a baseline for the F18 dataset, similar to the approach taken with the B24 dataset in the main paper. Peak push-off force (PO) predictions within the F18 dataset using *Model2* are shown in Fig. A4. Note the difference in sample sizes: F18 Young includes 24 participants, while F18 Elderly has only 10. Both predictions, from Young to Elderly (MAE = 0.15 N/kg, MAPE = 12.8%) and from Elderly to Young (MAE = 0.14 N/kg, MAPE = 14.1%) perform comparably well. As indicated in Table A3, F18 Elderly does not benefit from cadence, whereas F18 Young does; however, there is little difference in prediction quality between *Model1* and *Model2* within the F18 dataset. Specifically, MAPE for Young predicting Elderly changes only slightly from 12.8% in *Model2* to 13.5% in *Model1*; similarly, MAPE for Elderly predicting Young shifts from 14.1% in *Model2* to 13.7% in *Model1*). This minor difference occurs because both models adjust their parameter coefficients when cadence is included or excluded (see Table A6, data for the same range of walking speeds).

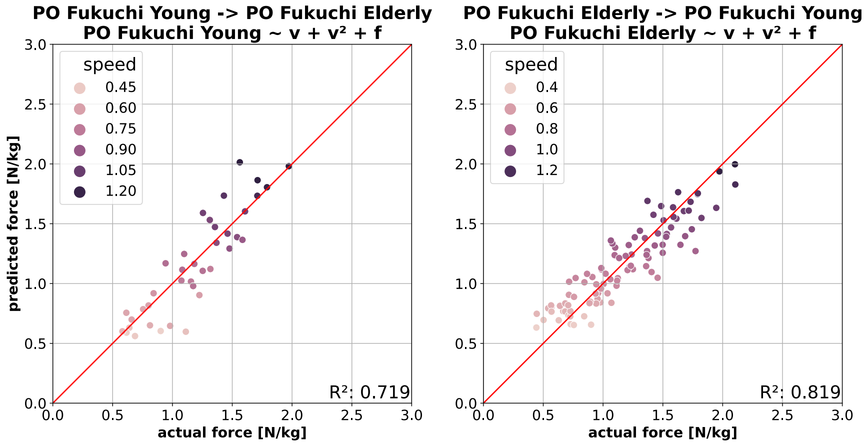


Figure A4: PO predictions within F18 dataset using *Model2*. Walking speeds are colour-coded. (***Left***) Model trained on Young data predicting PO from Elderly. (***Right***) Model trained on Elderly data predicting PO from Young.

Cross-dataset predictions – using *Model2* trained on B24 dataset to predict F18, and vice versa – are shown in Fig. A5. Models trained on B24 *Free Left* data (top row) consistently overestimate PO for both F18 Young and Elderly regardless of the speed, with similar performance: MAE = 0.21 N/kg and MAPE = 22.3% when predicting Young, and MAE = 0.21 N/kg and MAPE = 19.1% when predicting Elderly. The difference in age between F18 Young and Elderly has no impact on PO prediction accuracy. Conversely, models trained on F18 data (bottom row) tend to underestimate B24 *Free Left* data at all speeds. Notably, the model trained on F18 Elderly performs less accurately as it does not account for cadence, as discussed earlier. Despite this, both models show similar prediction error: the model trained on F18 Young predicts with MAE = 0.24 N/kg and MAPE = 18.8%, while the model trained on F18 Elderly predicts with MAE = 0.26 N/kg and MAPE = 18.5%.

The influence of including/excluding cadence is consistent across model estimation for both F18 and B24 datasets, as shown in Table A6. Adding cadence to *Model1* increases $v$ and decreases $v^2$ coefficient, regardless of whether only the preferred or all cadences are used in training B24 models. Notably, though, when predicting F18 from the B24 dataset using *Model1* (without cadence), pre-

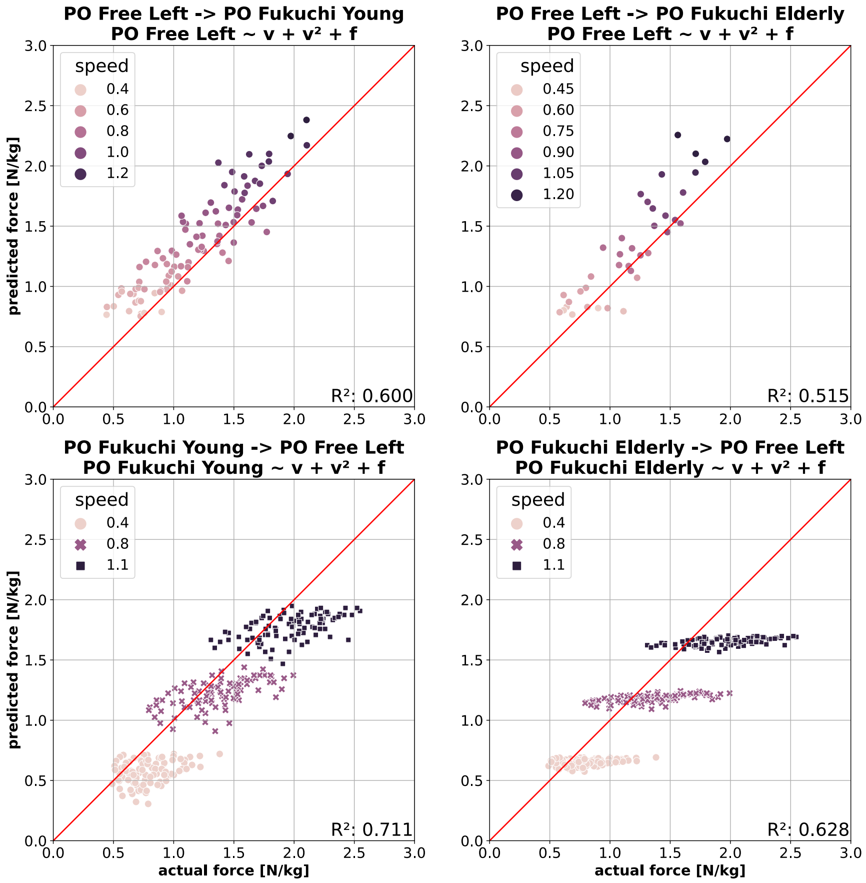


Figure A5: Cross-dataset predictions of PO. (***Top***) Peak push-off force (PO) *Model2* trained on B24 *Free Left* data predicting F18 Young and Elderly data. (***Bottom***) Peak push-off force (PO) *Model2* trained on F18 Young or Elderly data predicting B24 *Free Left* data.

Table A6: Model parameter coefficients in B24 and F18 datasets - *Model1* vs. *Model2*.

| | B24 Free Left | | | | F18 Young | | F18 Elderly | |
|---|---|---|---|---|---|---|---|---|
| | Model1 *(pref f)* | Model2 *(pref f)* | Model1 *(all f)* | Model2 *(all f)* | Model1 | Model2 | Model1 | Model2 |
| **speed (v)** | 0.588 | 1.204 | 0.658 | 1.483 | 1.364 | 2.191 | 0.894 | 1.175 |
| **speed (v²)** | 0.723 | 0.562 | 0.680 | 0.451 | 0.206 | 0.001 | 0.361 | 0.263 |
| **cadence** | / | -0.007 | / | -0.008 | / | -0.008 | / | -0.002 |
| **intercept** | 0.444 | 0.646 | 0.417 | 0.675 | -0.024 | 0.233 | 0.210 | 0.290 |

*B24 tested with all 5 cadences per speed (“all f”), and only with preferred cadence per speed (“pref f”).*

diction accuracy for F18 Elderly declines substantially: $R^2$ drops from 0.515 (as shown in Fig. A5) to 0.254 (not visualised), while $R^2$ for predicting F18 Young remains largely unchanged, decreasing only from 0.600 (as shown in Fig. A5) to 0.552 (not visualised). This difference is also reflected in prediction errors: when predicting F18 Young, the accuracy of the model trained on B24 changes in MAE from 0.21 N/kg to 0.23 N/kg and MAPE from 22.3% to 23.9%. This is a smaller decline in prediction accuracy compared to predicting F18 Elderly, where MAE declines from 0.21 N/kg to 0.28 N/kg and MAPE from 19.1% to 25.8%.

#### Step length

We begin by establishing a baseline for step length (SL) predictions within the F18 dataset using *Model1* (see Fig. A6). Note that the number of data points differs between graph due to difference in participant counts between the Young and Elderly datasets. Both predictions – from Young to Elderly (MAE = 0.04 N/kg, MAPE = 9.30%) and from Elderly to Young (MAE = 0.04 N/kg, MAPE = 8.11%) – are comparably accurate. Prediction quality remains very similar when the full range of speeds is used as well (not visualised here), with MAE unchanged and MAPE slightly reduced to 8.25% for Young predicting Elderly and to 7.28% for Elderly predicting Young.

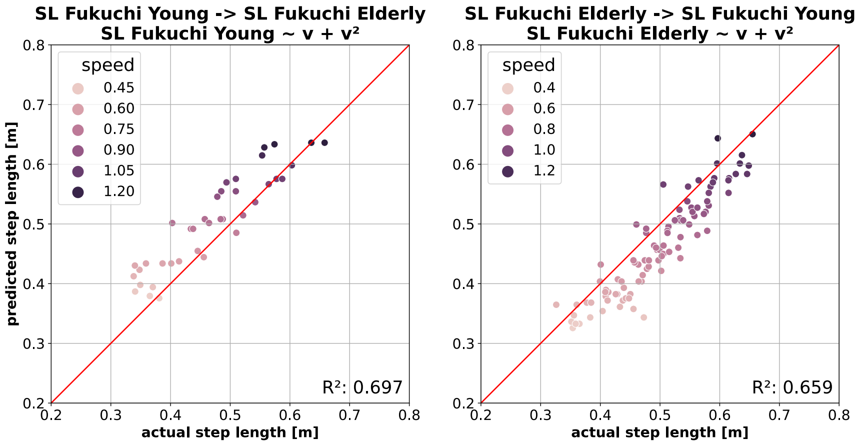


Figure A6: SL predictions within F18 dataset using *Model1*. Walking speeds are colour-coded. (***Left***) Model trained on Young data predicting SL from Elderly. (***Right***) Model trained on Elderly data predicting SL from Young.

Cross-dataset predictions – using *Model1* trained on the B24 dataset to predict F18 data – are shown in Fig. A7. Models trained on B24 *Free Left* data predict F18 Young with considerably higher accuracy than F18 Elderly, as indicated by a significant difference in $R^2$ and lower prediction errors for Young (MAE = 0.03 N/kg vs. 0.04 N/kg, and MAPE = 4.7% vs. 9.9%, Young vs. Elderly). Across speeds, the B24-trained model tends to overestimate step length for F18 Elderly, consistent with the inter-dataset predictions within the F18 dataset shown in Fig. A6 (left).

To investigate whether Age makes a difference in model prediction quality, we trained models on both B24 and F18 datasets by adding Age as an input model parameter to *Model1*. The results of cross-dataset predictions using these models are visualised in Fig. A8. Adding age does not alter prediction accuracy of the B24-trained model in predicting F18 Young (compare Fig. A7, left and Fig. A8, top left): $R^2$ and MAE (0.22 N/kg) remain the same, while MAPE changes from 6.4% to 4.7%, respectively. Conversely, adding Age to the B24-trained model makes predicting F18 Elderly

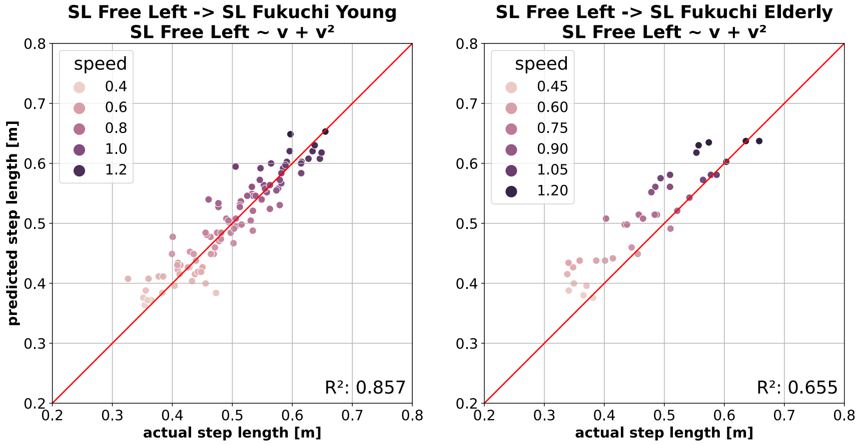


Figure A7: Cross-dataset predictions of SL. The model is trained on B24 *Free Left* data using speed only (*Model1*). (***Left***) Prediction of SL from F18 Young. (***Right***) Prediction of SL from F18 Elderly.

worse, with $R^2$ reducing from 0.655 to 0.513, MAE from 0.04 N/kg to 0.05 N/kg, and MAPE from 9.8% to 12.1% (compare Fig. A7, right and Fig. A8, top right).

A prediction of SL from the B24 dataset using models trained on the F18 dataset (*Model1* extended with *Age*) is visualised in Fig. A8 (bottom row). Only data from walking at the preferred cadence in B24 are included. As shown, both predictions (F18 Young and Elderly) achieve comparable accuracy, with $R^2$ values indicating strong predictive performance. This is further supported by the MAE (0.05 N/kg for both Young and Elderly) and MAPE (11.7% for Young and 12.3% for Elderly).

Prediction accuracy remains consistent when F18 Young *Model1* (without *Age*) is used to predict B24: $R^2$ and MAE remain virtually unchanged, and MAPE shows only a slight improvement (11.1%). These results indicate that F18 Young can predict B24 SL effectively with or without *Age* as a model parameter, mirroring the performance of B24 when predicting F18 Young. However, the same is not true when predicting B24 using F18 Elderly *Model1*. In this case, predictive accuracy as measured by $R^2$ drops significantly from 0.810 (Fig. A8, bottom right) to 0.645 (not visualized), while MAE increases to 0.06 N/kg and MAPE to 12.2%.

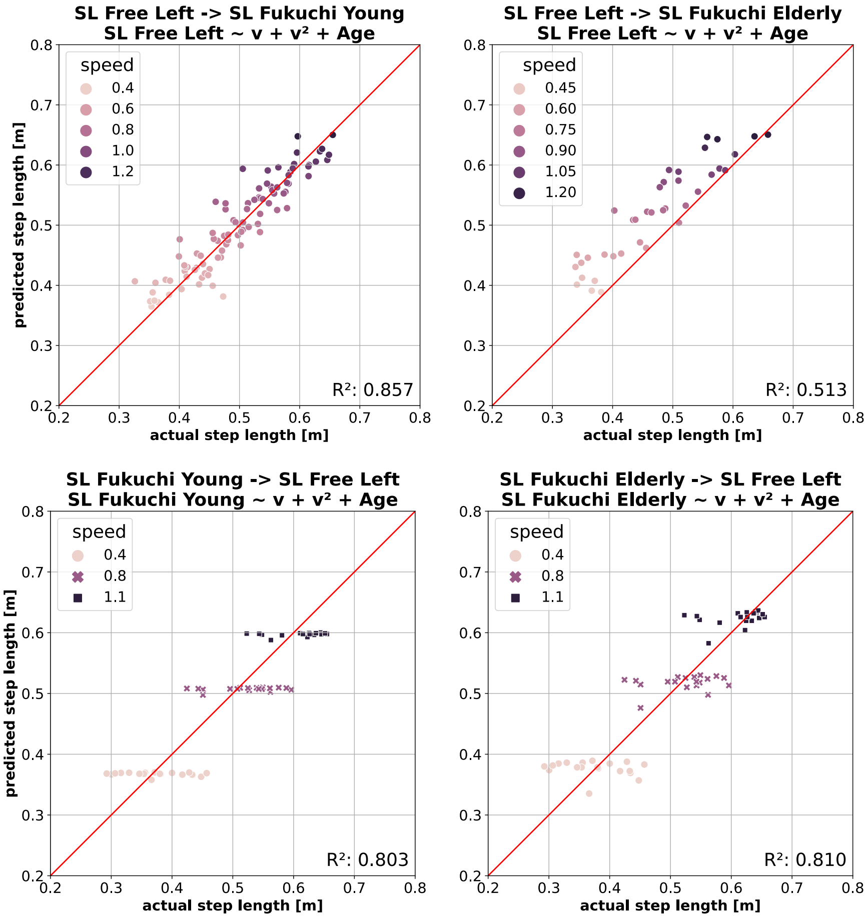


Figure A8: Cross-dataset predictions of SL using *Model1* (speed only) extended by adding Age. (***Top***) Prediction of SL from F18 Young (*left*) and Elderly (*right*) using model trained on B24 data. (***Right***) Prediction of SL from B24 using model trained on F18 Young (*left*) and F18 Elderly (*right*).
